# Single-cell analyses reveal SARS-CoV-2 interference with intrinsic immune response in the human gut

**DOI:** 10.1101/2020.10.21.348854

**Authors:** Sergio Triana, Camila Metz Zumaran, Carlos Ramirez, Carmon Kee, Patricio Doldan, Mohammed Shahraz, Daniel Schraivogel, Andreas R. Gschwind, Lars M. Steinmetz, Carl Herrmann, Theodore Alexandrov, Steeve Boulant, Megan L. Stanifer

## Abstract

**Objective:** Exacerbated pro-inflammatory immune response contributes to COVID-19 pathology. Despite the evidence about SARS-CoV-2 infecting the human gut, little is known about the importance of the enteric phase of SARS-CoV-2 for the viral lifecycle and for the development of COVID-19-associated pathologies. Similarly, it remains unknown whether the innate immune response triggered in this organ to combat viral infection is similar or distinct compared to the one triggered in other organs.

**Design:** We exploited human ileum- and colon-derived organoids as a non-transformed culture model supporting SARS-CoV-2 infection. We characterized the replication kinetics of SARS-CoV-2 in intestinal epithelial cells and correlated the expression of the viral receptor ACE2 with infection. We performed conventional and targeted single-cell transcriptomics and multiplex single-molecule RNA fluorescence in situ hybridization and used IFN-reporter bioassays to characterize the response of primary human intestinal epithelial cells to SARS-CoV-2 infection.

**Results:** We identified a subpopulation of enterocytes as the prime target of SARS-CoV-2. We found the lack of positive correlation between susceptibility to infection and the expression of ACE2 and revealed that SARS-CoV-2 downregulates ACE2 expression upon infection. Infected cells activated strong proinflammatory programs and produced interferon, while expression of interferon-stimulated genes was limited to bystander cells due to SARS-CoV-2 suppressing the autocrine action of interferon in infected cells.

**Conclusion:** Our findings reveal that SARS-CoV-2 curtails the immune response in primary human intestinal epithelial cells to promote its replication and spread and this highlights the gut as a proinflammatory reservoir that should be considered to fully understand SARS-CoV-2 pathogenesis.

**Significance of the study:** **What is already known about this subject?**

- COVID-19 patients have gastrointestinal symptoms which likely correlates with SARS-CoV-2 infection of the intestinal epithelium
- SARS-CoV-2 replicates in human intestinal epithelial cells.
- Intestinal organoids are a good model to study SARS-CoV-2 infection of the gastrointestinal tract
- There is a limited interferon response in human lung epithelial cells upon SARS-CoV-2 infection.

**What are the new findings?**

- A specific subpopulation of enterocytes are the prime targets of SARS-CoV-2 infection of the human gut.
- There is a lack of correlation between ACE2 expression and susceptibility to SARS-CoV-2 infection. SARS-CoV-2 downregulates ACE2 expression upon infection.
- Human intestinal epithelium cells produce interferon upon SARS-CoV-2 infection.
- Interferon acts in a paracrine manner to induce interferon stimulated genes that control viral infection only in bystander cells.
- SARS-CoV-2 actively blocks interferon signaling in infected cells.

**How might it impact on clinical practice in the foreseeable future?**

- The absence of correlation between ACE2 levels and susceptibility suggest that medications influencing ACE2 levels (e.g. high blood pressure drugs) will not make patients more susceptible to SARS-CoV-2 infection.
- The restricted cell tropism and the distinct immune response mounted by the GI tract, suggests that specific cellular restriction/replication factors and organ specific intrinsic innate immune pathways can represent unique therapeutic targets to treat COVD-19 patients by considering which organ is most infected/impacted by SARS-CoV-2.
- The strong pro-inflammatory signal mounted by the intestinal epithelium can fuel the systemic inflammation observed in COVID-19 patients and is likely participating in the lung specific pathology.

## Introduction

Coronavirus Disease 2019 (COVID-19) is caused by the severe acute respiratory syndrome coronavirus 2 (SARS-CoV-2). This highly infectious zoonotic virus has caused a global pandemic with almost 30,000,000 people infected worldwide as of September 2020. An exacerbated pro-inflammatory immune response generated by the host has been proposed to be responsible for the symptoms observed in patients [1–3]. Numerous studies have correlated the nature and extent of the immune response with the severity of the disease [4–6]. While many countries have succeeded in curtailing the first wave of infection, there is growing evidence that a second wave of infection is expected to take place and has even already started in some countries. Therefore, it is very urgent that we understand the virus-induced pathogenesis, in particular the immune response generated by the host, to develop prophylactic therapeutics, antiviral approaches and pharmacological strategies to control and revert the pathologies seen in patients.

SARS-CoV-2 is a member of the betacoronavirus genus, which initiates its lifecycle by exploiting the cellular receptor angiotensin converting enzyme 2 (ACE2) to enter and infect host cells [7]. Virus entry relies not only on ACE2, but also on the cellular proteases furin and the transmembrane serine protease 2 (TMPRSS2) that cleave and activate the SARS-CoV-2 envelope spike protein [8]. Following release of the genome into the cytosol, translation of the positive strand RNA genome is initiated and viral proteins quickly induce the formation of cellular membrane-derived compartments for virus replication and *de novo* assembly of virus particles. The host cells execute several strategies to counteract viral replication. Cellular pathogen recognition receptors (PRRs) sense viral molecular signatures or pathogen-associated molecular patterns (PAMPs) and induce a signaling cascade leading to the induction of interferons (IFNs) and pro-inflammatory molecules. IFNs represent the first line of defense against viral infection as their autocrine and paracrine signaling leads to the production of hundreds of interferon-stimulated genes (ISGs) known to exert broad antiviral functions [9].

SARS-CoV-2 infection is not limited to the respiratory tract and COVID-19 patients show systemic manifestation of the disease in multiple organs [10,11]. For many of these organs, it is unclear whether the pathology is a side effect of SARS-CoV-2 infection in the lung and its associated pro-inflammatory response or whether it is due to a direct SARS-CoV-2 infection of the specific organ. For the gastrointestinal (GI) tract, there is clear evidence of SARS-CoV-2 replication which is associated with the release of viral genome into the feces [12–15]. Human intestinal organoids have been established as a robust model to study SARS-CoV-2 infection and provided direct evidence about primary human intestinal epithelial cells efficiently supporting SARS-CoV-2 replication [16–18]. Importantly, while SARS-CoV-2 infection in the lung is characterized by a curtailed IFN response [19,20], the intrinsic immune response in intestinal epithelial cells is characterized by the production of IFN and ISGs, with IFNs providing some protection to intestinal epithelium cells against SARS-CoV-2 [17]. Studies in human intestinal organoids revealed that only discrete cells are susceptible to SARS-CoV-2 infection and some evidence suggests that these cells may be enterocytes [16]. However, the precise cell tropism of SARS-CoV-2 within the colon and other parts of the gastrointestinal tract is yet to be fully characterized. Finally, despite the driving role of inflammation in the pathologies observed in COVID-19 patients, we are still lacking important molecular details concerning the inflammatory response generated by SARS-CoV-2 infected cells and how the surrounding bystander cells will respond to it.

Here, we aim to address the outlined gaps by applying single-cell RNA-sequencing to human ileum- and colon-derived organoids infected with SARS-CoV-2. Using differential gene expression analysis and multiplex single-molecule RNA fluorescence *in situ* hybridization (FISH), we investigated the cell type tropism of SARS-CoV-2 and its link to ACE2 expression levels. While we could show that immature enterocytes represent the primary site of SARS-CoV-2 infection, we did not observe correlation between infectivity and ACE2 expression. Interestingly, we could observe that SARS-CoV-2 infection is associated with a downregulation of ACE2 expression. Pathway analysis revealed that infected cells mount a strong pro-inflammatory response characterized by the upregulation of both NFκB/TNF expression and the activity of their respective pathways. On the contrary, bystander cells were characterized by an upregulation of the IFN-mediated immune response as monitored by the increased production of ISGs. Importantly, using a combination of multiplex single molecule RNA FISH and IFN-reporter bioassays we could show that while IFN could act in a paracrine manner in bystander cells, IFN cannot act in an autocrine manner in SARS-CoV-2 infected cells. Our findings demonstrate that SARS-CoV-2 has developed strategies to impair IFN-mediated signaling in infected cells, and together with our previous observations showing that IFN restricts SARS-CoV-2 replication in intestinal cells [17], these results suggest that SARS-CoV-2 manipulates the cell intrinsic innate immune response to promote its replication and spread.

## Results

### Single-cell RNA sequencing of SARS-CoV-2 infected colon and ileum organoids

A fraction of COVID-19 patients show enteric symptoms and it has been shown that SARS-CoV-2 replicates in the intestinal tract of patients [13] and in human primary intestinal epithelial cells [16–18,21]. To characterize SARS-CoV-2 interactions with primary human intestinal epithelial cells (hIECs) and to ultimately address whether the enteric phase of SARS-CoV-2 contributes to the observed cytokine profiles and systemic inflammation observed in COVID-19 patients, human intestinal organoids were infected by SARS-CoV-2. To address whether organoids derived from distinct parts of the intestinal tract display different susceptibility, colon- and ileum-derived organoids were infected by SARS-CoV-2 and their response was monitored along the course of infection. For both colon- and ileum-derived organoids, we could observe the presence of infected cells as early as 4 hours post-infection (hpi) with the number of infected cells increasing within the course of infection (Fig. S1A). This was corroborated with an increase in intracellular viral RNA and the release of *de novo* infectious virus particles over time (Fig. S1B-C) thus demonstrating efficient virus replication and spreading of infection in both colon and ileum organoids. To characterize how hIECs respond to SARS-CoV-2 infection, we monitored the production of type I IFN (IFNβ1) and type III IFNs (IFNλ1 and IFNλ2-3) over time. SARS-CoV-2 did not induce significant production of IFNβ1 in either ileum or colon organoids, except for a slight upregulation of IFNβ1 expression in colon organoids at 24 hpi (Fig. S1D, left panels). On the contrary, for IFNλ2-3, a strong upregulation was observed in both colon and ileum organoids upon infection by SARS-CoV-2 (Fig. S1D, right panels). Interestingly, similar to IFNβ1, IFNλ1 was not upregulated in response to infection (Fig. S1D, middle panels). Taken together, these results show that a fraction (around 10%) of human intestinal epithelial cells fully supports SARS-CoV-2 infection, replication, *de novo* production of infectious virus particles and that infection is associated with the upregulation of type III IFNs (IFNλ2-3) (Fig. S1A).

### Determination of SARS-CoV-2 intestinal epithelial cell tropism

Human intestinal organoids are composed of multiple cell types recapitulating the cellular complexity of the human intestinal epithelium. Although it is clear that SARS-CoV-2 infects the human intestinal epithelium, which specific cell types are infected by the virus, how infection impacts the transcriptional landscape of each individual cell type and how bystander cells respond to viral infection remains unknown. To characterize SARS-CoV-2 interactions with hIECs at the single-cell transcriptional level, colon- and ileum-derived organoids were infected with SARS-CoV-2 and subjected to single-cell RNA sequencing (scRNAseq) (Fig. 1A). Single-cell suspensions were generated and 3’ scRNAseq was performed across six biological conditions (mock, 12 hpi and 24 hpi for both colon and ileum organoids). scRNAseq provided broad transcriptional profiles for around 2000 cells per condition with 5000 and 6000 genes profiled on average per cell for the colon and the ileum, respectively (Fig. S2A-F for colon organoids and Fig. S2G-L for ileum organoids).

**Figure 1.**
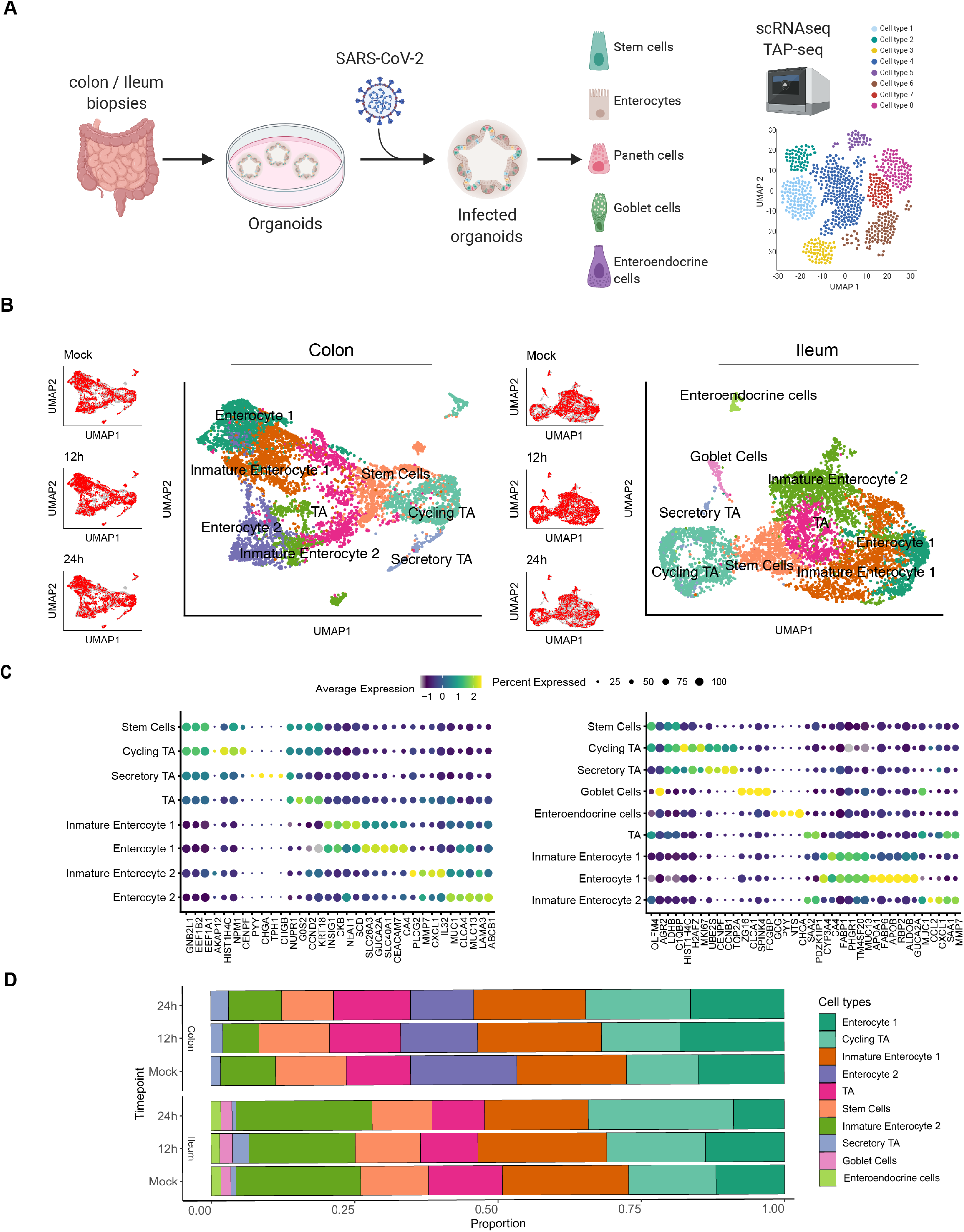
Single cell sequencing of SARS-CoV-2 infected colon- and ileum-derived human organoids. A. Schematic representation of the experimental workflow. B. Uniform manifold approximation and projection (UMAP) embedding of single-cell RNA-Seq data from mock and SARS-CoV-2 infected colon-derived (left panels) and ileum-derived (right panels) organoids colored according to the cell type. Small insets represent the UMAP for mock and infected organoids at 12 and 24 hpi. C. Dot plot of the top marker genes for each cell type for (left) colon- and (right) ileum-derived organoids. The dot size represents the percentage of cells expressing the gene; the color represents the average expression across the cell type. D. Bar plot displaying the proportion of each cell type in mock and infected organoids (12 and 24 hpi).

To identify the cell types present in our organoids, we used the Uniform Manifold Approximation and Projection (UMAP) algorithm for dimensionality reduction of our scRNAseq data. We identified eight and nine clusters of cells in the colon and ileum organoids, respectively (Fig. 1B). Using both cell-type-specific marker genes (Fig. 1C) from single-cell atlases of human intestinal tissues [22]and our annotated scRNAseq data from human Ileum biopsies [23], we could identify the cell types represented in these clusters (Fig. 1B). Stem cells, transient amplifying (TA) cells, enterocytes, goblet and enteroendocrine cells were found to be present in intestinal organoids (Fig. 1B). Different subpopulations of enterocytes in ileum and colon (CLCA4+, ALDOB+, MUC13+), were identified, namely, enterocytes 1 (GUCA2A+, FABP6+, CA4+), enterocytes 2 (MMP7+,MUC1+, CXCL1+) as well as immature enterocytes 1 and immature enterocytes 2, the latter likely representing cells not fully differentiated into mature enterocytes. The presence of two distinct populations of enterocytes and their immature related forms is consistent with previous reports [22]. Importantly, infection by SARS-CoV-2 did not alter cell clustering (Fig. 1B, left UMAP inset panels) and did not impact the proportions of the different cell types present in both the colon and ileum organoids (Fig. 1D) thus allowing us to investigate cell tropism.

To increase the sensitivity and dynamic range in detecting the SARS-CoV-2 genome and selected host genes, we made use of additional targeted scRNA-seq (Schraivogel et al. 2020) performed on the same organoid samples. Targeted scRNA-seq is more sensitive to detect genes-of-interest irrespective of their expression level, and quantifies gene expression with a higher dynamic range compared to conventional 3’ scRNAseq [24]. We selected 12 genes, including the SARS-CoV-2 genome, the SARS-CoV-2 receptor ACE2, an interferon stimulated gene (ISG15) and a panel of hIEC type markers that we previously determined by scRNAseq of ileum biopsies and organoids (APOA4, CHGB, FABP6, FCGBP, LYZ, MKI67, OLFM4, SLC2A2, SMOC2 and SST) [23]. Looking at the relative expression of SARS-CoV-2 genome in mock vs infected cells (Fig. S3A-B), classical 3’ scRNAseq detected the viral genome in a proportion of organoid cells (Fig. S3A (inset); ~75% at 12 hpi and ~25% at 24 hpi), while using the targeted approach, SARS-CoV-2 counts were detected in virtually all cells in the infected samples (Fig. S3B). Concurrently, we observed that the number of SARS-CoV-2 counts per cell increased over the course of infection both targeted and whole transcriptome scRNA-seq (Fig. S3A-B), consistent with active replication of the virus in organoids monitored using q-RT-PCR (Fig. S1B). Since it is expected that only a fraction of the cells present in intestinal organoids are indeed infected by SARS-CoV-2 (Fig. S1A) [16,17], the presence of SARS-CoV-2 genome in all cells likely does not reflect active viral replication, but could be explained by the presence of viruses attached to the cells surface or by free-floating viral particles or RNA. Capitalizing on the high dynamic range provided by targeted scRNA-seq, we defined cells having productive infection and replication as those with the SARS-CoV-2 counts higher than a baseline, calculated as the mean expression of SARS-CoV-2 measured by targeted scRNA-seq in all cells-containing droplets (Fig. S3C-D, top panels). Cells with targeted scRNA-seq SARS-CoV-2 expression levels below the estimated baseline level were defined as non-infected bystander cells. Following this approach, we could correct for the presence of contaminating viral RNA. Only a small fraction of cells (Colon: 4.5% and Ileum: 5.2%) was detected as supporting SARS-CoV-2 infection (Fig. S3C-D, UMAP bottom panels) which is consistent with our immunofluorescence staining of SARS-CoV-2 infected organoids (Fig. S1A) and other reports [16,17].

Comparing the targeted scRNA-seq and the conventional 3’ scRNAseq approaches, we observed high correlation between the expression levels estimated by these technologies (Fig. S3E-F top panels), with targeted scRNA-seq providing several orders of magnitude higher dynamic range, compared to the conventional 3’ scRNA-seq (Fig. S3E-F bottom panels). The high positive correlation between the two sequencing approaches (except for SLCA2) serves as a quality control confirming that targeted scRNAseq could robustly quantify both viral and cellular genes.

While SARS-CoV-2 could be detected in most cell types in both colon and ileum organoids (Fig. 2A), immature enterocytes 2 consistently represented the main virus-targeted cell type (Fig. 2B). The proportion of infected immature enterocytes 2 and the number of SARS-CoV-2 genome copy per cell increased over the course of infection (Fig. 2B) which is consistent with both the increasing number of infected cells observed by immunofluorescence and the increasing copies of SARS-CoV-2 genome over time (Fig. S1A-B, S3A-B). In ileum-derived organoids, TA cells in the secretory lineage were also found to be highly infected by SARS-CoV-2 (Fig. 2B, right panel). Interestingly, these cells were infected mostly at 24 hpi, suggesting that they are secondary targets of infection. Taken together, these results suggest that immature enterocytes 2 are the primary target of SARS-CoV-2 infection in hIECs both in colon and ileum.

**Figure 2.**
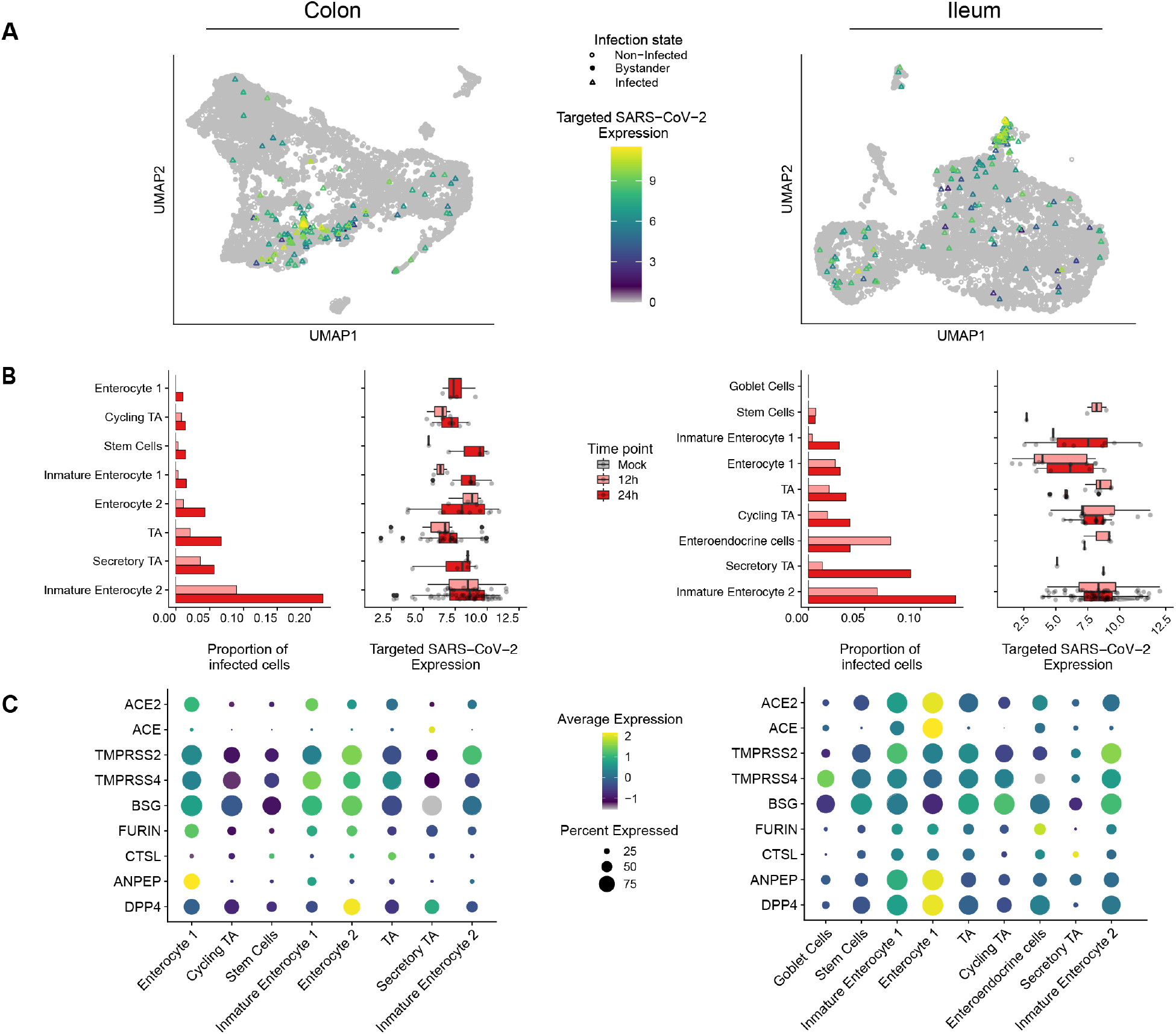
SARS-CoV-2 cell tropism in human colon- and ileum-derived organoids. A. UMAP visualization of scRNAseq data of SARS-CoV-2 infected colon-(left) and ileum-derived organoids (right). Triangles represent infected cells and colors represent the corrected targeted normalized expression of SARS-CoV-2 determined using the targeted scRNAseq data. B. Proportion of cells infected with SARS-CoV-2 for each cell type and corresponding normalized expression values of SARS-CoV-2 in each individual cell type in colon (left) and ileum (right) organoids. Data are color coded for mock, 12 and 24 hpi. C. Dot plots of known entry determinants across cell types. The dot size represents the percentage of cells expressing the gene; the color represents the average expression across the cell type. Data are from mock-infected colon (left) and ileum organoids (right).

### SARS-CoV-2 cell tropism and association with expression of ACE2 and TMPRSS2

The angiotensin-converting enzyme 2 (ACE2) and the cellular protease type II transmembrane serine protease 2 (TMPRSS2) are known to be major determinants for SARS-CoV-2 infection. ACE2 is the cellular receptor of SARS-CoV-2 mediating viral entry [7]. TMPRSS2 is a cellular protease that processes the SARS-CoV-2 spike (S) protein which is an essential step for viral envelope fusion with the host membrane and release of viral contents in the cytosol of the cells. Combined conventional and targeted scRNAseq enabled us to investigate the link between SARS-CoV-2 genome copy numbers and expression of ACE2 in a cell type-specific manner. Different to what we have expected, immature enterocytes 2, the main site of SARS-CoV-2 infection in both colon and ileum organoids (Fig. 2A-B), were not the cells displaying the highest levels of ACE2 (Fig. S4A-B). Analysis of ACE2 expression levels in all cell types revealed that cells with relatively high levels of ACE2 (*e.g.* enterocytes 1) were not susceptible to SARS-CoV-2 infection (Fig. 2B-C). Similarly, we found that SARS-CoV-2 infection is not associated with the expression of the receptor structural homologue ACE, a candidate receptor for SARS-CoV-2 basigin (BSG, also known as CD147), as well as the cellular proteases furin, Cathepsin L1 (CTSL), aminopeptidase ANPEP and DPP4 (MERS-CoV receptors) (Fig. 2C). On the contrary, TMPRSS2 was found to be highly expressed in immature enterocytes 2 (Fig. 2C). In summary, although ACE2 is a recognized receptor for SARS-CoV-2, we found no association between ACE2 expression levels and susceptibility to infection on the single-cell level or across detected types of hIECs.

### SARS-CoV-2 infection induces downregulation of ACE2 expression in intestinal organoids

ACE2 has been reported to act as an interferon stimulated gene (ISG) resulting in an increased expression level upon viral infection and interferon stimulation of nasal and lung epithelial cells [25]. Similarly, in COVID-19 patients, ACE2 expression was shown to be upregulated in lung epithelial cells compared to control patients [26]. To investigate whether the expression of either ACE2 or other putative receptors and key cellular proteases is upregulated upon SARS-CoV-2 infection or upon SARS-CoV-2-mediated immune response in primary hIECs, we compared their expression levels in mock-infected cells *vs.* both SARS-CoV-2-infected and non-infected bystander cells. Although we did not observe any association between ACE2 expression in non-infected cells and their propensity to be infected by SARS-CoV-2 (Fig. 2B-C and S4A-B), differential gene expression analysis revealed that upon SARS-CoV-2 infection the ACE2 expression levels were downregulated (Fig. 3A-B). In colon organoids, visible downregulation of ACE2 expression was observed in infected cells, progressing from 12 hpi to 24 hpi, as compared to mock-infected cells (Fig. 3B, left panel). Importantly, no significant difference of ACE2 expression in the bystander cells was observed (Fig. 3B). In ileum-derived organoids, ACE2 expression was also downregulated in the infected cells. However, in contrast to colon organoids, ACE2 expression in bystander cells of ileum organoids was also downregulated as compared to mock-infected cells (Fig. 3B, right panels). Moreover, ACE2 expression was found to be negatively correlated with the presence of the viral genome (Fig. 3C-D). The downregulation of ACE2 expression was not only observed in immature enterocytes 2 which were identified as the primary site of SARS-CoV-2 infection (Fig. 2B), but it was also observed in most other cell types present in ileum-derived organoids over the course of SARS-CoV-2 infection (Fig. 3E). Expression levels of the other SARS-CoV-2 putative receptors and of key cellular proteases (e.g. TMPRSS2, furin and CTSL) were also found to be reduced in both infected and bystander cells in ileum derived-organoids as compared to mock-infected cells (Fig. 3B, right panel). Interestingly, when considering colon-derived organoids, the expression levels of these cellular genes were found slightly increased at 12 hpi and decreased at 24 hpi (Fig. 3B, left panel). Altogether, these data suggest that ACE2 expression is downregulated in colon- and ileum-derived hIECs upon SARS-CoV-2 infection. To validate this observation, we performed multiplex single-molecule fluorescence *in situ* hybridization (FISH) on SARS-CoV-2 infected organoids. At 12 and 24 hpi, organoids were fixed and evaluated using transcript-specific probes directed against the SARS-CoV-2 genome and ACE2. Fluorescence microscopy analysis confirmed that infected cells indeed display lower expression levels of ACE2 at both 12 hpi and 24 hpi (Fig. 3F, white arrow). Quantification of the relative expression levels of SARS-CoV-2 genome and ACE2 transcripts in the RNA FISH images at the single-cell level again confirmed a negative correlation between SARS-COV-2 and ACE2 (Fig. 3G). Altogether, our data strongly suggest that the expression levels of ACE2 decrease in both colon and ileum hIECs upon SARS-CoV-2 infection.

**Figure 3.**
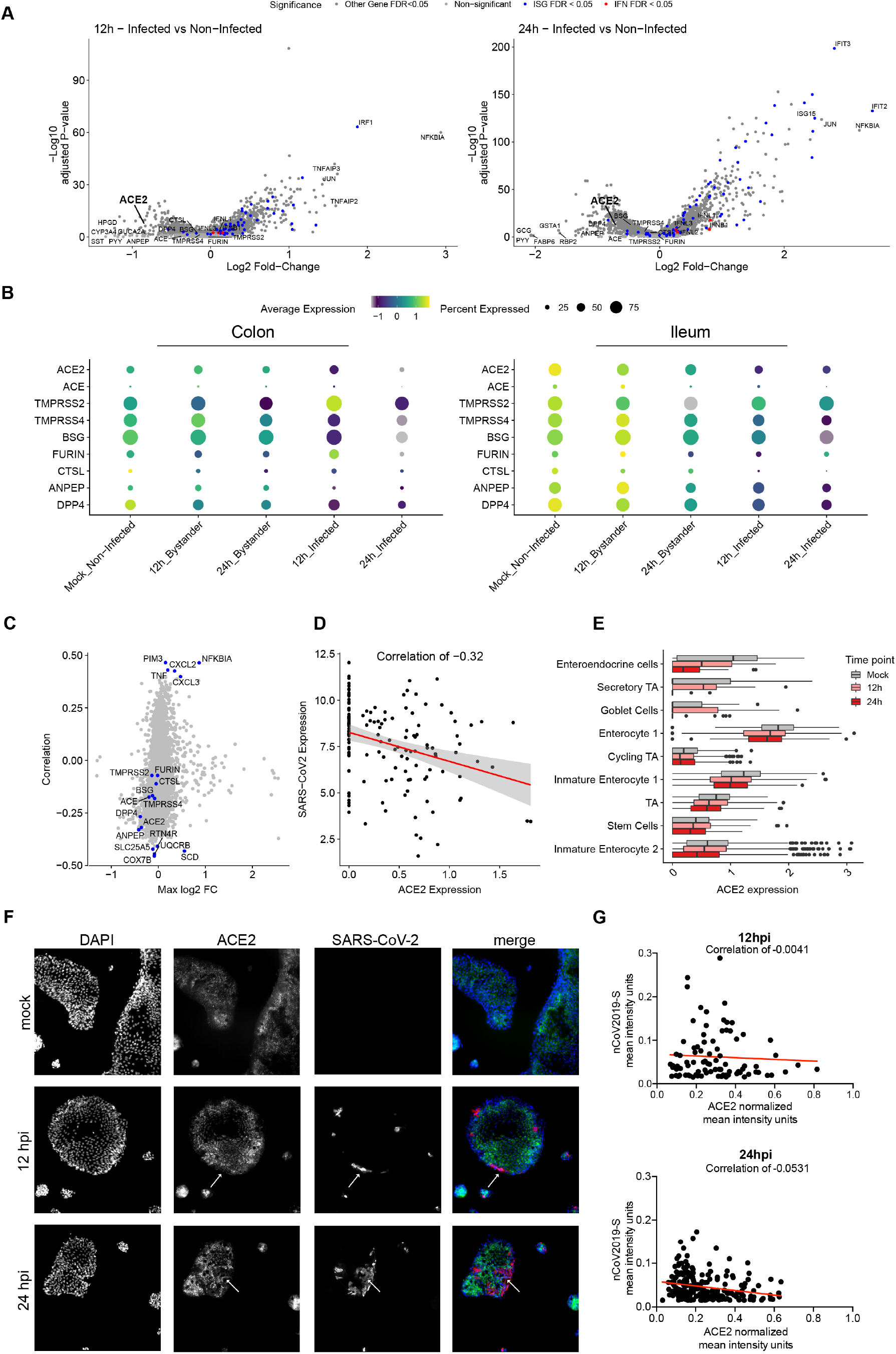
Downregulation of ACE2 upon SARS-CoV-2 infection. A. Volcano plots of genes that are differentially expressed in infected cells relative to mock-infected cells at 12 hpi and 24 hpi in ileum organoids. The statistical significance (−log10 of the adjusted p-value) is shown as a function of the log2 fold change. B. Dot plots displaying the expression changes of known SARS-CoV-2 entry determinants for both infected and bystander cells during the course of infection (mock, 12 and 24 hpi) in colon (left) and ileum organoids (right). The dot size represents the percentage of cells expressing the gene; the color represents the average expression across the cell type excluding zeros. C. Correlation of gene expression values with the amount of SARS-CoV-2 genome (y-axis) vs the maximal log2 fold change (x-axis) across conditions. This plot is generated by comparing both 12 and 24 hpi to mock. D. Correlation of SARS-CoV-2 expression with ACE2 expression across all cell types at 24 hpi. E. Expression values of ACE2 in each cell type for mock infected and SARS-CoV-2 infected cells in ileum organoids at 12 hpi and 24 hpi. F. Multiplex in situ RNA hybridization of ACE2 and SARS-CoV-2 of mock-infected and infected 2D ileum organoids at 12 hpi and 24 hpi. White arrows point at SARS-CoV-2-infected cells. Representative image is shown. G. Correlation of the relative expression SARS-CoV-2 and ACE2 for infected cells from the multiplex in situ RNA hybridization shown in F. Each dot represents a cell.

### SARS-CoV-2 induces a pro-inflammatory response in hIECs

To evaluate the response of hIECs to SARS-CoV-2 infection, we performed a comparative gene expression analysis between mock-infected and infected organoids. For the infected organoids, we considered separately the infected cells (those with SARS-CoV-2 genome detected) and the bystander cells (those without SARS-CoV-2 genome). In colon organoids, already at 12 hpi hIECs display a strong NFκB and TNF response to infection with this response becoming even more pronounced at 24 hpi (Fig. S5A-B). When comparing mock to bystander cells, we noticed that at 24 hpi, the response of bystander cells mostly followed an IFN-mediated immune response characterized by the presence of multiple ISGs (Fig. S5C-D). This observation suggests that infected cells generate a pro-inflammatory response while bystander cells likely respond to the secreted IFN in a paracrine manner. This is supported by the differential gene expression analysis of bystander vs. infected cells showing that infected cells have a stronger NFκB and TNF mediated response compared to bystander cells (Fig. S5E-F). Similar results were found in SARS-CoV-2 infected ileum-derived organoids (Fig. S5G-L). Interestingly, at 24 hpi, some interferon-stimulated genes (ISGs) (e.g. IFIT1-3, MX1, CXCL10, IRF1) were found to be also upregulated in infected cells but to a much lesser extent compared to bystander cells (Fig. S5G-J). Additionally, while infected cells in ileum-derived organoids were found to generate a similar NFκB/TNF-mediated response compared to the colon-derived organoids (Fig. S5B and H), ileum-derived bystander cells had a stronger IFN-mediated response which can be seen by the overall higher expression of ISGs in ileum organoids compared to colon organoids upon SARS-CoV-2 infection (Fig. S5D and J). Together, these comparative gene expression analyses revealed that upon SARS-CoV-2 infection of human intestinal epithelial cells, both strong pro-inflammatory and IFN-mediated responses are generated.

### Cell type specific immune response in infected vs. bystander cells

Taking into account the differences in the susceptibility of the different hIEC types to SARS-CoV-2, with immature enterocytes 2 constituting the main site of SARS-CoV-2 infection (Fig. 2), we compared the response of each individual cell type to SARS-CoV-2 infection. Similar to the analysis of all cell types taken together (Fig. S5), differential gene expression analysis of colon-derived infected immature enterocytes 2 revealed a strong NFκB/TNF-mediated response while bystander immature enterocytes 2 mostly display an IFN-mediated response (Fig. 4A-C and Fig. S6A). Similarly, in ileum-derived organoids, infected immature enterocytes 2 also showed a strong NFκB/TNF-mediated response (Fig. 4F, 4H and S6B) while bystander cells were characterized by their response to secreted IFNs leading to ISG expression (Fig. 4G and Fig. S6B)

**Figure 4.**
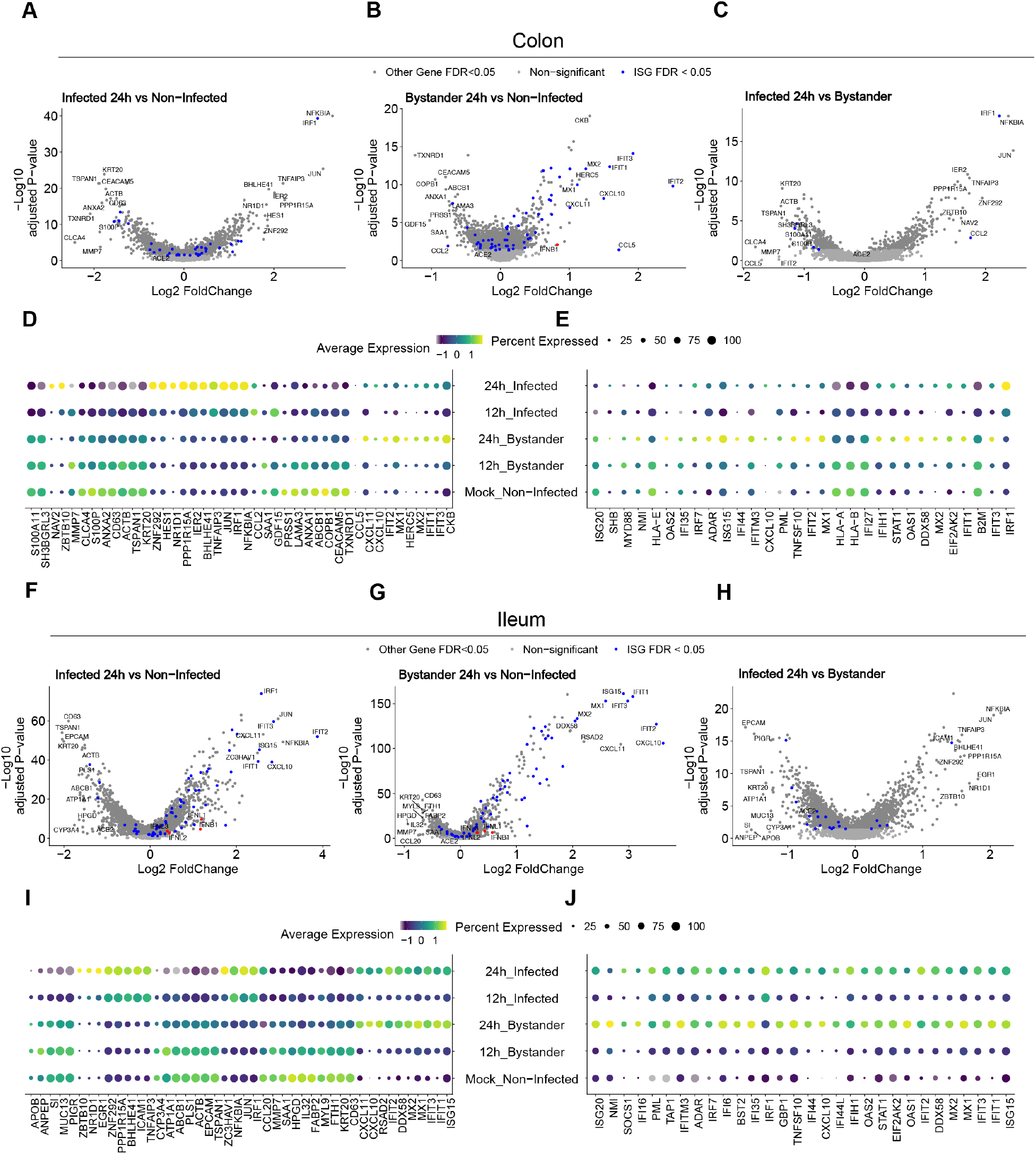
Intrinsic innate immune response generated by immature enterocytes 2 upon SARS-CoV-2 infection. (A-C) Volcano plots displaying the genes that are differentially expressed in immature enterocytes 2 upon SARS-CoV-2 infection of colon organoids. A. infected vs. mock infected cells, B. bystander vs. mock infected cells and C. infected vs. bystander cells. The statistical significance (−log10 adjusted p-value) is shown as a function of the log2 fold change. D. Dot plot of the top 42 most differentially expressed genes upon SARS-CoV-2 infection in mock, infected and bystander cells at 12 and 24 hpi. The dot size represents the percentage of cells expressing the gene; the color represents the average expression across the cell type. E. Same as in D but for the top 30 most differentially expressed ISGs. F-G. Same as A-E but for ileum organoids.

Pathway analysis confirmed that the bystander response was mostly an IFN-related response while the infected cell response was mostly pro-inflammatory (Fig. S7). Comparison of the transcriptional response to SARS-CoV-2 infection in infected *vs.* bystander immature enterocytes 2 further confirmed that infected cells mount a strong pro-inflammatory response characterized by the upregulation of NFκB and TNF (Fig. 4C-D and 4H-I). Analysis of the top 30 differentially expressed ISGs in colon-derived organoids clearly shows that at 24 hpi, bystanders cells respond to IFN by upregulating the expression of a large panel of ISGs (Fig. 4E). Similar findings were observed in immature enterocytes 2 from ileum organoids, although infected cells were also found to express more ISGs compared to their colon-derived counterparts. Importantly, in both colon and ileum organoids, bystanders showed higher levels of expression for all considered ISGs compared to infected cells (Fig. 4E and 4J). These results are consistent with the observation that ileum-derived organoids are more immune-responsive compared to colon-derived organoids (Fig. S5D and J). Together these data show that upon SARS-CoV-2 infection, infected cells mount a strong NFκB/TNF-mediated pro-inflammatory response and display a limited production of ISGs while bystander cells mount a strong IFN-mediated response through a strongly-upregulated expression of a broad panel of ISGs.

To determine if the characterized NFκB/TNF-high and IFN-low immune response is specific to immature enterocytes 2, we also looked individually at each infected cell type. We found that in all considered types of hIECs, bystander cells display an IFN-mediated response during SARS-CoV-2 infection while infected cells are characterized by a strong NFκB/TNF-mediated response (Fig. S8-9). Since our targeted scRNAseq analysis revealed the presence of background viral RNA in the samples (Fig. S3), we asked whether the observed immune response is indeed cascading to bystander cells through type III interferon secreted by infected cells or is caused by the direct action of non-replicating viral particles on bystander cells. To address this, colon and ileum organoids were infected with either live or UV-inactivated SARS-CoV-2. Results revealed that upon infection with live SARS-CoV-2 both IFNs and ISGs were produced (Fig. S10). On the contrary, exposure of organoids to UV-inactivated SARS-CoV-2 did not lead to virus replication and cells failed to produce both IFN and ISGs (Fig. S10). This demonstrates that active replication is required for the described immune response and allowed us to rule out the exposure to non-replicating viral particles as being the cause of this response. Altogether our results show that infected and bystander cells respond differently to SARS-CoV-2 infection where infected cells mount a NFκB/TNF-mediated response while bystander cells mount a IFN-mediated response.

### Signaling activity in infected vs. bystander cells

To characterize the signaling that underpins the distinct immune response of infected and bystander cells upon SARS-CoV-2 infection, we inferred the pathway signaling activity from scRNAseq data with PROGENy (Fig. 5A-B). For both colon and ileum organoids, infected cells show a strong activation of the MAPKs, NFκB and TNFα pathways. In line with the enrichment analysis (Fig. S7, S8), these pathways were found to be less activated in bystander cells with higher scores in ileum compared to colon (Fig. 5A-B). Interestingly and in accordance with our differential gene expression analyses (Fig. 4, S5 and S6), the JAK-STAT signalling pathway was found to be activated mostly in bystander cells (Fig. 5A-B). To further elucidate the signalling activity at the single-cell level, we generated diffusion maps of all single cells based on the scRNAseq expression of interferon-related genes (Fig. 5C-D). In both iluem and colon, we observed a clear bifurcation of all cells into two distinct branches, one branch representing mainly infected cells and another branch representing mainly bystander cells. In addition, we calculated the activities of selected transcription factors (TFs) for all single cells using SCENIC and mapped the inferred activities onto the single-cell diffusion maps (Fig. 5C-D, right insets). We found that the transcription factors STAT1 and IRF1 were activated mainly in bystander cells (branch along DC1) while JUN was activated in infected cells (branch along DC2) (Fig. 5C-D, left panels). Extending this analysis to transcription factors whose activity pattern is highly correlated to either DC1 or DC2 revealed that globally, transcription factors that are critical for IFN-mediated signaling (i.e. the ISGF3 complex: STAT1/STAT2/IRF9 and IRF1) are highly active in bystander cells (Fig. 5E-F). Similarly, the ETS variant transcription factor 7 (ETV7) which is an ISG acting as a negative regulator of IFN-mediated signaling [27] and the Early Growth Response Gene 1 (EGR1) which enhances type I IFN signalling [28] were also found to be activated in bystander cells (Fig. 5E-F). Upregulation of the EGR1 and JUN dependent pathways was consistent with the findings of the previous work investigating SARS-CoV-2 infection of the human lung and intestinal epithelial cell lines Calu-3 and Caco-2 [29].

**Figure 5.**
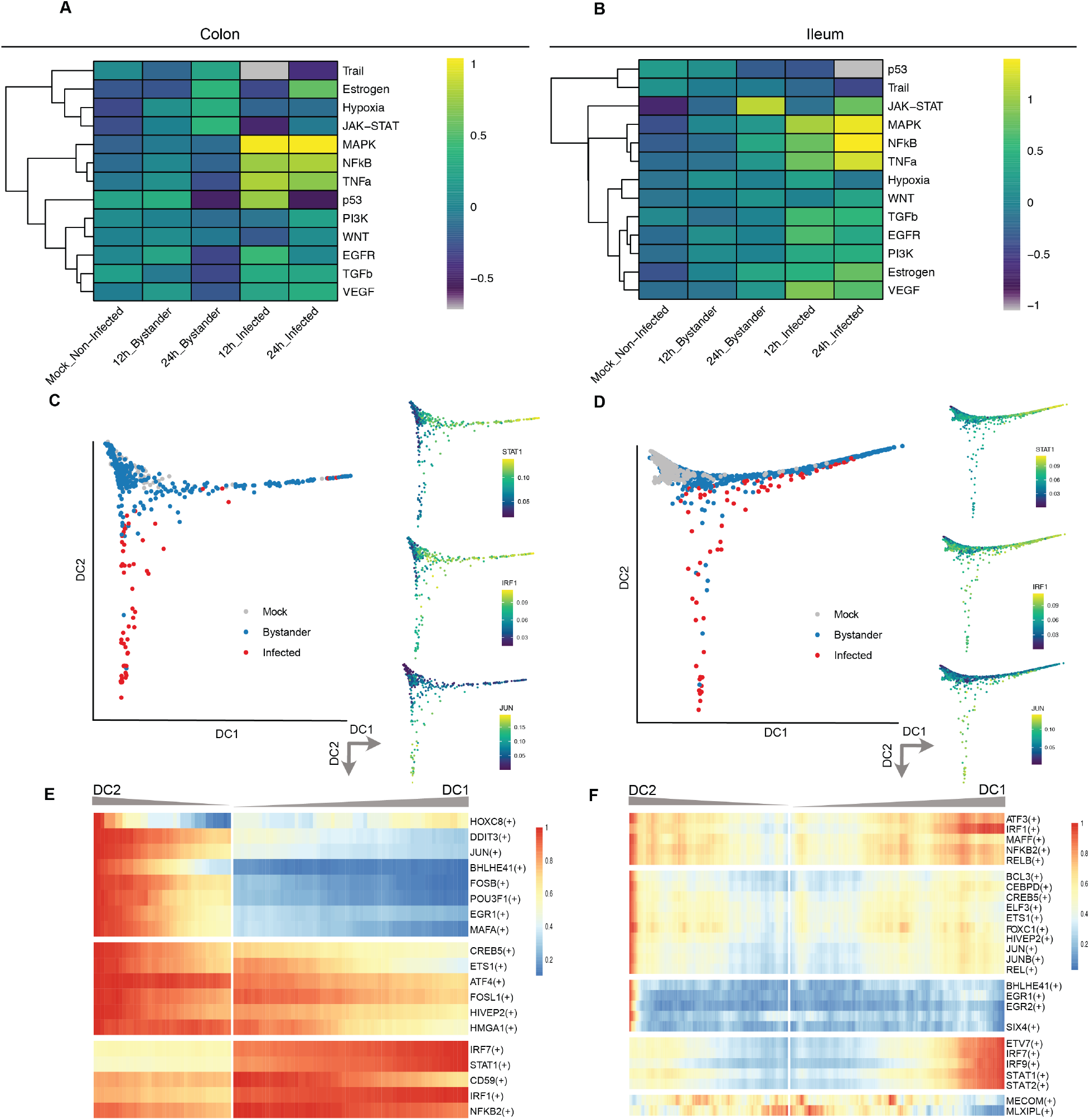
Differential signaling activity of infected vs. bystander cells upon SARS-CoV-2 infection. A-B. Heatmaps of pathway signaling activity inferred by PROGENy for A. colon and B. ileum organoids. C-D. Diffusion maps embeddings showing mock, bystander and infected cells (big panels) and activities for selected transcription factors STAT1, IRF1 and JUN (small panels) for C. colon and D. ileum organoids. E-F. Heatmaps of transcription factor activities along bifurcating trajectories of the corresponding diffusion maps for E. colon and F. ileum organoids. The dimensions DC1 and DC2 represent the first two eigenvectors of the Markovian transition matrix and were calculated separately for either colon or ileum organoids.

### SARS-CoV-2 inhibits IFN-mediated ISG expression

To validate that the IFN-mediated response is specific to bystander cells, ileum-derived organoids were infected with SARS-CoV-2. At 12 and 24 hpi, single molecule RNA FISH was performed using probes specific for the SARS-CoV-2 genome and for ISG15 which was found to be highly upregulated upon infection and has the highest −log10 *p*-value in the differential analysis comparing bystander cells vs mock-infected cells (Fig. 4G). Microscopy images revealed that bystander cells (non-infected) were indeed positive for ISG15 (Fig. 6A). Interestingly, SARS-CoV-2 infected cells were found to express little to no ISG15 (Fig 6A, white arrows). Quantification confirmed that cells which displayed the highest expression of ISG15 were negative for SARS-CoV-2 genome (Fig. 6B). These results support the model that bystander cells respond to SARS-CoV-2 infection by mounting an IFN-mediated response. On the other hand, as shown by using both scRNAseq (Fig. 4) and RNA FISH (Fig. 6A), infected cells do not mount an IFN-mediated response (low-to-no expression of ISGs) suggesting that infected cells are refractory to IFN. To address whether SARS-CoV-2 infection can indeed impair IFN-mediated signaling, we directly monitored ISG production in response to IFN treatment in either infected or non-infected cells. However, while it is well known that ISGs are made in a JAK-STAT dependent manner downstream the IFN receptors, there is growing evidence that a subset of ISGs can be made downstream IRF3 following viral recognition by PRRs. To alleviate this unwanted IRF3 mediated production of ISGs, we generated a human intestinal cell line (T84) depleted of IRF3. Infection of the IRF3 KO T84 did not result in the production of ISGs as monitored by q-RT-PCR of ISG15 (Fig. 6D). IRF3 KO T84 cells were mock infected or infected with SARS-CoV-2, at 24 hpi cells were treated with IFN and 12 h post-treatment production of ISG15 was assessed by q-RT-PCR (Fig. 6C). Results show that mock infected IRF3 KO T84 cells were responsive to IFN demonstrating that genetic depletion of IRF3 does not alter IFN-mediated signaling (Fig. 6D). Interestingly, in SARS-CoV-2 infected IRF3 KO T84 cells, production of ISG15 upon IFN treatment was significantly downregulated (Fig. 6D). To confirm that this impaired induction of ISG15 upon IFN treatment was specific to SARS-CoV-2 infection, IRF3 KO T84 cells were infected with astrovirus at an MOI of 3 to achieve full infection (Fig. 6E). Contrary to SARS-CoV-2 infection, infection of IRF3 KO T84 cells by astrovirus did not impair IFN-mediated signaling as a similar upregulation of ISG15 was observed in both mock-infected and astrovirus-infected cells upon IFN treatment (Fig. 6D). In a second validation approach, to fully demonstrate that only infected cells have an altered IFN-mediated signaling, we developed an assay based on a fluorescent reporter of ISG expression (Fig. 6F). For this, we generated a T84 cell line transduced with a reporter made of the promoter region of the ISG MX1 driving the expression of the fluorescent protein mCherry. Mx1 is known to be made strictly downstream of the IFN receptor in a JAK-STAT dependent manner but not downstream of IRF3. Upon IFN treatment, about 30-40% of cells expressing this reporter were responsive and became fluorescent (Fig. 6G). Following infection with SARS-CoV-2 at an multiplicity of infection (MOI) of 3, most of the cells were found to be infected (Fig. 6G). However when cells were treated 24 hpi with IFN, most infected cells did not respond to IFN and very few became double positive for both virus and MX1 driven mCherry (Fig. 6G, left panel). To control that non-infected bystander cells could indeed respond to IFN and express mCherry, we repeated this experiment but using an MOI of 1 for SARS-CoV-2 infection. About 40% of the cells were found to be infected. Supplementing IFN affected mainly non-infected cells, as can be seen from the increase of MX1-positive cells and no change in the number of double-positive cells (both infected and MX1-positive) (Fig. 6G, right panel).

**Figure 6.**
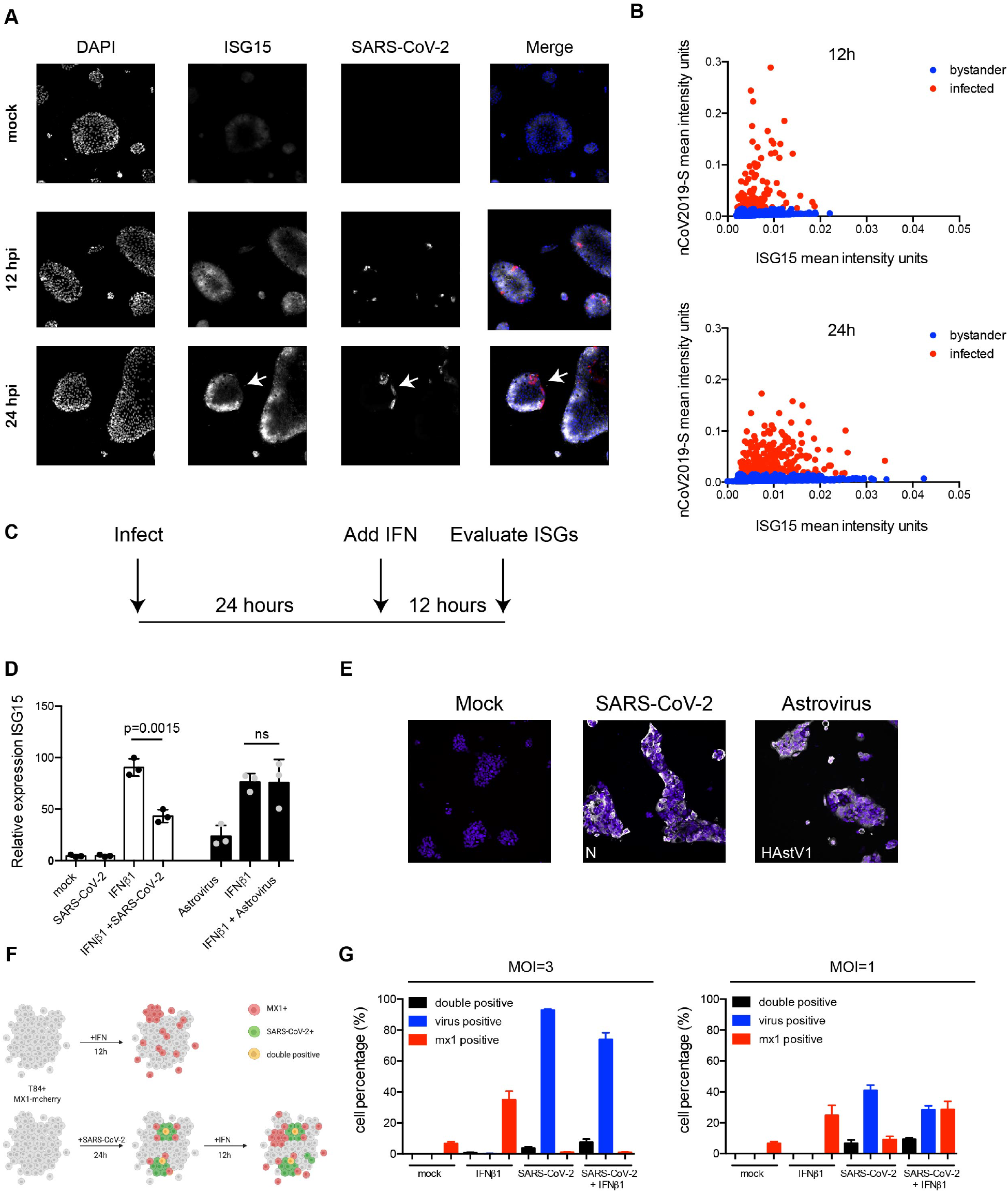
SARS-CoV-2 infection impairs interferon-mediated signaling. A. Human ileum-derived organoids were seeded in 2D on iBIDi chamber slides. 12 and 24 hpi cells were fixed and the amount of SARS-CoV-2 infected cells (red) and the induction of ISG15 (white) was analyzed by single molecule RNA FISH. Nuclei were visualized with DAPI (blue). Representative image is shown. B. Quantification of the mean fluorescence intensity of samples in A. Each dot represents a cell. C. Schematic depicting the infection and interferon addition to evaluate ISG15 induction following SARS-CoV-2 and astrovirus infection. D. T84 IRF3 knock-out cells were infected with SARS-CoV-2 or human astrovirus 1 at an MOI=3. 24 hpi, cells were incubated in the presence or absence of 2000 IU/mL of IFNβ1. 12 h post-treatment RNA was harvested and the induction of ISG15 was analyzed by q-RT-PCR. Three biological replicates were performed. Error bar indicates standard deviation. Statistics were determined by unpaired t-test. E. T84 IRF3 knock-out cells were infected with SARS-CoV-2 or human astrovirus 1 at an MOI=3. 24 hpi, cells were fixed and stained for either the SARS-CoV-2 N protein or for human astrovirus 1 (HAstV1) capsid protein. Experiments were performed in triplicate. Representative image is shown. F. Schematic depicting the T84 MX-1 mcherry expressing cells and their response to IFN or SARS-CoV-2 infection. G. T84 MX-1 mcherry were infected with SARS-CoV-2 at an MOI=3 or MOI=1. 24 hpi cells were incubated in the presence or absence of 2000 IU/mL of IFNβ1. 12 h post-treatment cells were fixed and stained for SARS-CoV-2 N protein. The number of MX-1 positive, SARS-CoV-2 positive and double positive cells was quantified. Three biological replicates were performed. Error bar indicates standard deviation.

**Figure 7.**
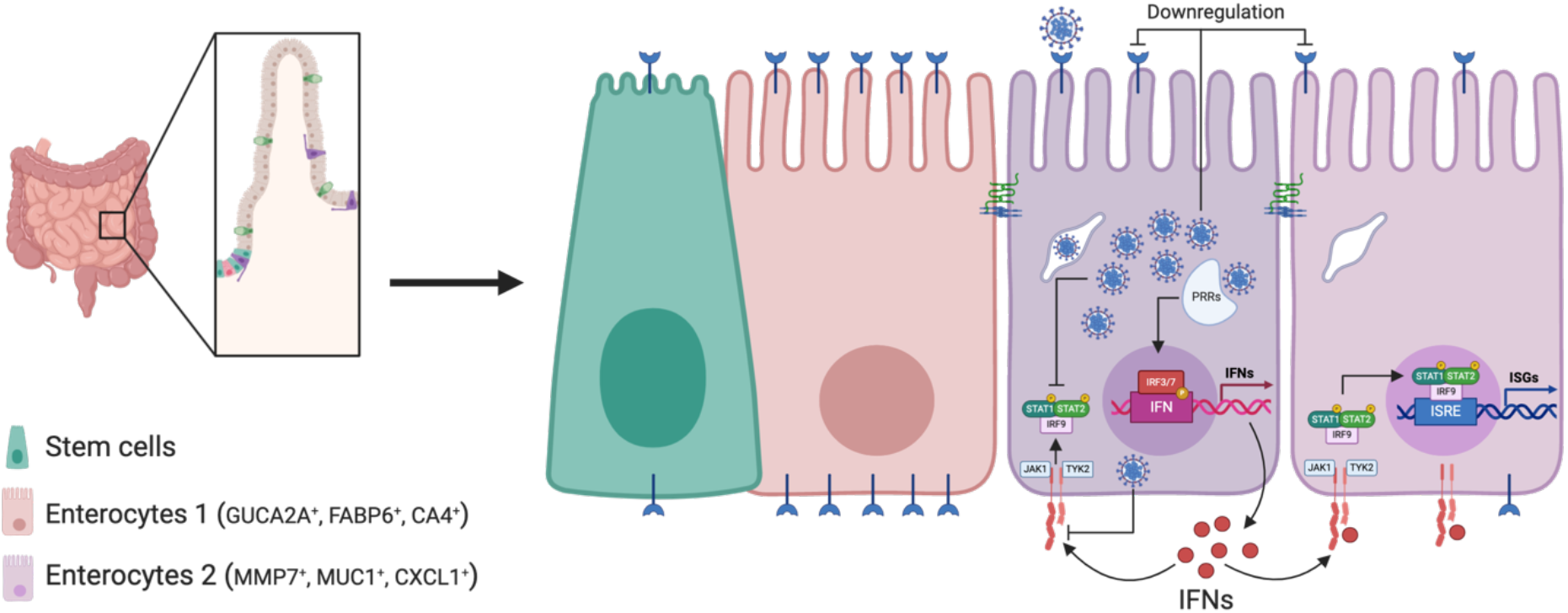
Schematic of SARS-CoV-2 infection of human intestinal epithelial cells. SARS-CoV-2 infects a subpopulation of enterocytes. Upon infection, enterocytes mount a pro-inflammatory response characterized by the upregulation of NFκB and TNF. Bystander cells respond to secreted IFN and upregulate the expression of ISGs. SARS-CoV-2 infection induces the downregulation of ACE2 expression and interferes with IFN-mediated signalling in infected cells.

Altogether, our results provide a strong evidence that upon infection with SARS-CoV-2 primary human intestinal cells generate a strong NFκB/TNF-mediated response and produce IFN. This IFN acts in a paracrine manner onto bystander cells that leads to the upregulation of IFN-stimulated genes. Importantly, SARS-CoV-2 infection renders infected cells refractory to IFN as they show little-to-no increase in the activity of the JAK-STAT signaling pathways and fail to upregulate IFN-stimulated gene expression.

## Discussion

Many COVID-19 patients display gastroenteritis symptoms and there is growing evidence that the intestinal epithelium can be infected by SARS-CoV-2. Whether the symptoms are associated with the direct replication of SARS-CoV-2 in the GI tract or are a consequence of the strong pro-inflammatory response seen in patients is unclear. Using human intestinal “mini-gut” organoids has already demonstrated that intestinal epithelium cells can support SARS-CoV-2 infection, replication and spread. However, which cell type is infected remains poorly defined [16–18]. By exploiting single-cell transcriptomics approaches (scRNAseq) and targeted scRNAseq, we identified that a subpopulation of enterocytes (namely, immature enterocytes 2) is the cell type most susceptible to SARS-CoV-2 infection. Interestingly, other cell types also supported infection by SARS-CoV-2 but to a much lesser extent (Fig. 2B). The characterized tropism of SARS-CoV-2 could be explained by either cell type-specific intrinsic differences rendering some cell type more permissive or due to an overrepresentation of cells of a particular cell type. In our colon-derived organoids, there were twice as many immature enterocytes 1 compared to immature enterocytes 2 and in ileum-derived organoids both enterocytes lineages were present in roughly equal numbers. This suggests that the SARS-CoV-2 cell tropism for immature enterocytes 2 is not due to a higher proportion of these cells in our organoids but due to intrinsic differences between immature enterocytes 2 and other epithelial cell lineages. Differential gene expression analysis between immature enterocytes 2 and the most similar other annotated cell type (immature enterocytes 1) does not highlight the presence or absence of obvious restriction/replication factors that could explain the observed tropism.

The expression levels of the SARS-CoV-2 receptor ACE2 were found to be higher in immature enterocytes 1 while immature enterocytes 2 express more of the key cellular protease TMPRSS2 (Fig. 2C and S4). Although expression of ACE2 is mandatory for infection [7], we noticed no correlation between ACE2 expression level and the copy numbers of SARS-CoV-2 genome in the cell (Fig. 2B-C)(Fig. S11). This observation is important as it raises a question about using ACE2 expression as the only basis for conjectures about infectability of cell types or even organs, the approach recently proposed in various SARS-CoV-2-related publications [30–37]. Our results highlight that the investigation of SARS-CoV-2 tropism requires biological validation of infection and should not be done solely based on analysis of transcriptional profiles of individual cells or tissues. Interestingly, we found that TMPRSS2 expression levels were well-associated with the SARS-CoV-2 genome copy numbers in human intestinal epithelial cells (Fig. 2C and S4) which is consistent with the observation that TMPRSS2 and the related protease TMPRSS4 are critical for infection of intestinal organoids [18]. As such it is tempting to speculate that TMPRSS2 plays a more important role in the SARS-CoV-2 cell tropism than ACE2, however more studies are required to fully explore this hypothesis.

Several studies have suggested that ACE2 is an interferon-stimulated gene and is induced upon SARS-CoV-2 infection [25,26]. This led to a speculation that upon infection and induction of pro-inflammatory responses, the ACE2 levels would increase thereby favoring SARS-CoV-2 infection. Our results show that, on the contrary, upon infection ACE2 levels decrease both in infected and bystander hIECs (Fig. 3)(Fig. S11). Interestingly, differences in the kinetics of ACE2 regulation were observed between colon- and ileum-derived organoids. In colon-derived organoids, a small increase in ACE2 expression was observed at early times post-infection (12 hpi) but at later time points (24 hpi) the overall expression of ACE2 (and other putative SARS-CoV-2 receptors and key cellular proteases) was decreased as compared to mock infected cells (Fig. 3B). In ileum organoids the expression of ACE2 was decreased over the time course of infection. The observed upregulation of ACE2 upon infection might be tissue-specific and time-dependent. However, recently it has been proposed that ACE2 does not behave as an ISG but instead a novel form of ACE2 (dACE2) is interferon-inducible [38]. dACE2 results from transcription initiation at an internal exon leading to the production of an alternative short version. Within our scRNAseq data we could not distinguish between the two forms and as such the observed temporal increase (in colon-derived organoids) could be due to the downregulation of ACE2 with the concomitant upregulation of dACE2.

The nature of the PRR responsible for sensing SARS-CoV-2 infection is yet to be determined but from recent work and previous work on SARS-CoV-1 and MERS it could be speculated that TLR3, RLRs and the STING pathways could be involved [39,40]. In our current work we could show that active virus replication is required to induce an IFN-mediated response as infection by UV-inactivated SARS-CoV-2 did not lead to IFN and ISG production (Fig. S10). Interestingly, when human intestinal epithelial cells are infected by a UV-inactivated reovirus, which is a virus whose genome is a dsRNA, an IFN-mediated response is induced [41]. As SARS-CoV-2 is single-stranded RNA virus and dsRNA intermediates will only occur during active replication, it is tempting to speculate that what is being sensed are these dsRNA replication intermediates.

SARS-CoV-2 infection is characterized by a strong pro-inflammatory response and this has been observed both in tissue-derived samples and *in vitro* using cell culture models [19]. This pro-inflammatory response is characterized by upregulation of the NFκB and TNF pathways. Our scRNAseq approach revealed that this pro-inflammatory response is specific to infected cells and that bystander cells do not show a strong pro-inflammatory response. These differences between infected and bystander cells were earlier observed for other cell types: infection of human bronchial epithelial cells (HBECs) also reveal that the pro-inflammatory response is biased toward infected cells and not bystander cells [31].

It was reported that infection of human lung epithelial cells by SARS-CoV-2 is characterized by a low to absent IFN response [19]. On the contrary, in human intestinal epithelium cells an IFN-mediated response is readily detectable and is characterized by both the production of IFN and ISGs [16,17]. Interestingly, upon infection with SARS-CoV-2, we observed the upregulation of IFNλ2-3 but we failed to observe a significant increase in IFNλ1 expression (Fig. S1D). This absence of IFNλ1 upregulation is not specific to SARS-CoV-2 but a particularity of intestinal organoids, as a similar IFNλ2-3 specific response was observed when intestinal organoids were infected with other enteric viruses [42,43]. Upregulation of IFN production upon SARS-CoV-2 infection of intestinal epithelial cells was found to be low but significant (Fig. 4) and this could raise the question whether this small production of IFN is sufficient to induce the production of ISGs in a paracrine manner. Previous work on the antiviral function in type III IFN in human intestinal epithelial cells revealed that although type III IFN protects epithelial cells against infection in a dose-dependent manner, very small amounts of IFN are required to induce ISGs and to provide an antiviral state to the cells [42]. The current work confirms this model as despite a moderated upregulation of IFN expression in infected cells, we observed a strong ISG upregulation in bystander cells.

When comparing the immune response generated by organoids derived from different sections of the GI tract, we observed that ileum organoids were more immunoresponsive compared to colon organoids. While the extent of the up-regulation of genes related to pro-inflammatory response was similar between colon and ileum (Fig. 4A, 4F), we observed that ileum organoids, particularly bystander cells produced significantly more ISGs compared to their colon counterparts (Fig. 4b, 4G). This compartmentalization of response along the GI tract is consistent with previous reports describing that different sections of the GI tract respond differently to microbial challenges [44]. Most importantly, our results reveal that production of ISGs is mostly restricted to bystander cells, while production of IFN is detected mostly in infected cells (Fig. 5, S5 and S6). These observed differences between infected and bystander cells were confirmed using single-molecule RNA FISH showing that production of ISG15 was clearly observed in bystander and, to a much lesser extent, in infected cells (Fig. 6). An important finding of this work is that infected cells not only fail to produce ISGs, they also become refractory to IFN (Fig. 6) (Fig. S11). When SARS-CoV-2-infected intestinal cells were treated with IFN, only bystander cells upregulated ISG while infected cells did not. This absence of ISG induction in infected cells suggests that SARS-CoV-2 has developed mechanisms to shutdown IFN-mediated signaling and the subsequent production of ISGs (Fig. S11). Preventing IFN-mediated signaling in infected cells would provide a replication advantage to SARS-CoV-2 as secreted IFN will not be able to act in an autocrine manner to induce ISGs which will curtail virus replication and *de novo* virus production. Although the SARS-CoV-2 viral protein responsible for blocking the IFN-mediated signaling is yet to be identified in our system, a recent report suggests that ORF6 could block IFN-mediated signaling by interfering with STAT1 nuclear translocation [45].

In conclusion, in this work we identified a subset of immature enterocytes as the primary site of infection of SARS-CoV-2 in ileum- and colon-derived human intestinal epithelial cells. We could show that upon infection, infected cells mount a strong pro-inflammatory response characterized by a strong activation of the NFκB/TNF pathways while bystander cells mount an IFN-mediated response (Fig. S11). This differential response between infected and bystander cells is due to an active block of IFN-signaling in infected cells (Fig. S11). Interestingly, recent work performing scRNAseq of SARS-CoV-2 infected HBECs revealed that infected cells were readily responding to secreted interferon and produced large amounts of ISGs [31]. This suggests that there are cell type-specific or tissue-specific regulations of interferon-mediated signaling during SARS-CoV-2 infection. This needs to be considered when studying replication and pathogenesis of SARS-CoV-2 in different organs as well as when developing therapies against COVID-19.

## Acknowledgments

This work was supported by research grants from the Deutsche Forschungsgemeinschaft (DFG): project numbers 415089553 (Heisenberg program), 240245660 (SFB1129), 278001972 (TRR186), and 272983813 (TRR179) to SB; 416072091 to MS. CMZ is supported by the SFB1129 (240245660), CR and CH are supported by the TRR179 (272983813). We also acknowledge funding from the Helmholtz International Graduate School for Cancer Research to CK, the German Academic Exchange Service (DAAD) (Research Grant 57440921) to PD, Darwin Trust of Edinburgh to ST, and the ERC Consolidator Grant METACELL (773089) to TA.

## Author Contributions

ST performed experiments, analyzed the single cell data and contributed to manuscript writing. CR analyzed the single cell data. CMZ, CK, PD, MS and MLS performed experiments. DS and ARG performed target scRNA0seq. LS and CH contributed to the concept of the study and critical discussions. SB, MLS and TA conceived experiments, interpreted results and wrote the manuscript. The final version of the manuscript was approved by all authors.

## Methods

### Cells

T84 human colon carcinoma cells (ATCC CCL-248) and their knock-out derivative clones were maintained in a 50:50 mixture of Dulbecco’s modified Eagle’s medium (DMEM) and F12 (GibCo) supplemented with 10% fetal bovine serum and 1% penicillin/streptomycin (Gibco). Vero E6 (ATCC CRL 1586) were maintained in DMEM supplemented with 10% fetal bovine serum and 1% penicillin/streptomycin. The mCherry-tagged Mx1 promoter plasmid was a kind gift from Ronald Dijkman (University of Bern), which was used to generate a T84 stable cell line via lentiviral transduction. Single clones were derived from this cell line and evaluated for their ability to respond to both type-I and type-III interferons.

### Viruses

SARS-CoV-2 (strain BavPat1) was obtained from the European Virology Archive. The virus was amplified in Vero E6 cells.

### Human organoid cultures and ethic approval

Human tissue was received from colon resection or ileum biopsies from the University Hospital Heidelberg. This study was carried out in accordance with the recommendations of the University Hospital Heidelberg with informed written consent from all subjects in accordance with the Declaration of Helsinki. All samples were received and maintained in an anonymized manner. The protocol was approved by the “Ethics commission of the University Hospital Heidelberg” under the protocol S-443/2017. Organoids were prepared following the original protocol described by [46]. In brief, stem cells containing crypts were isolated following 2 mM EDTA dissociation of tissue samples for 1 h at 4°C. Crypts were spun and washed in ice cold PBS. Fractions enriched in crypts were filtered with 70 mM filters and the fractions were observed under a light microscope. Fractions containing the highest number of crypts were pooled and spun again. The supernatant was removed, and crypts were re-suspended in Matrigel. Crypts were passaged and maintained in basal and differentiation culture media as previously described [43].

### 2D organoid seeding

8-well iBIDI glass bottom chambers were coated with 2.5% human collagen in water for 1 h prior to organoids seeding. Organoids were collected at a ratio of 100 organoids/well. Collected organoids were spun at 450 xg for 5 mins and the supernatant was removed. Organoids were washed 1X with cold PBS and spun at 450 xg for 5 mins. PBS was removed and organoids were digested with 0.5% Trypsin-EDTA (Life technologies) for 5 mins at 37°C. Digestion was stopped by addition of serum containing medium. Organoids were spun at 450 xg for 5 mins and the supernatant was removed and organoids were re-suspended in normal growth media at a ratio of 250 μL media/well. The collagen mixture was removed from the iBIDI chambers and 250 μL of organoids were added to each well.

### Viral infections

Media was removed from cells and 10^6^ pfu of SARS-CoV-2 (as determined in Vero cells) was added to cells for 1 hour at 37°C. Virus was removed, cells were washed 1x with PBS and fresh media was added back to the cells.

### RNA isolation, cDNA, and qPCR

RNA was harvested from cells using RNAeasy RNA extraction kit (Qiagen) as per manufacturer’s instructions. cDNA was made using iSCRIPT reverse transcriptase (BioRad) from 250 ng of total RNA as per manufacturer's instructions. q-PCR was performed using iTaq SYBR green (BioRad) as per manufacturer’s instructions, TBP was used as normalizing genes. Primers used:

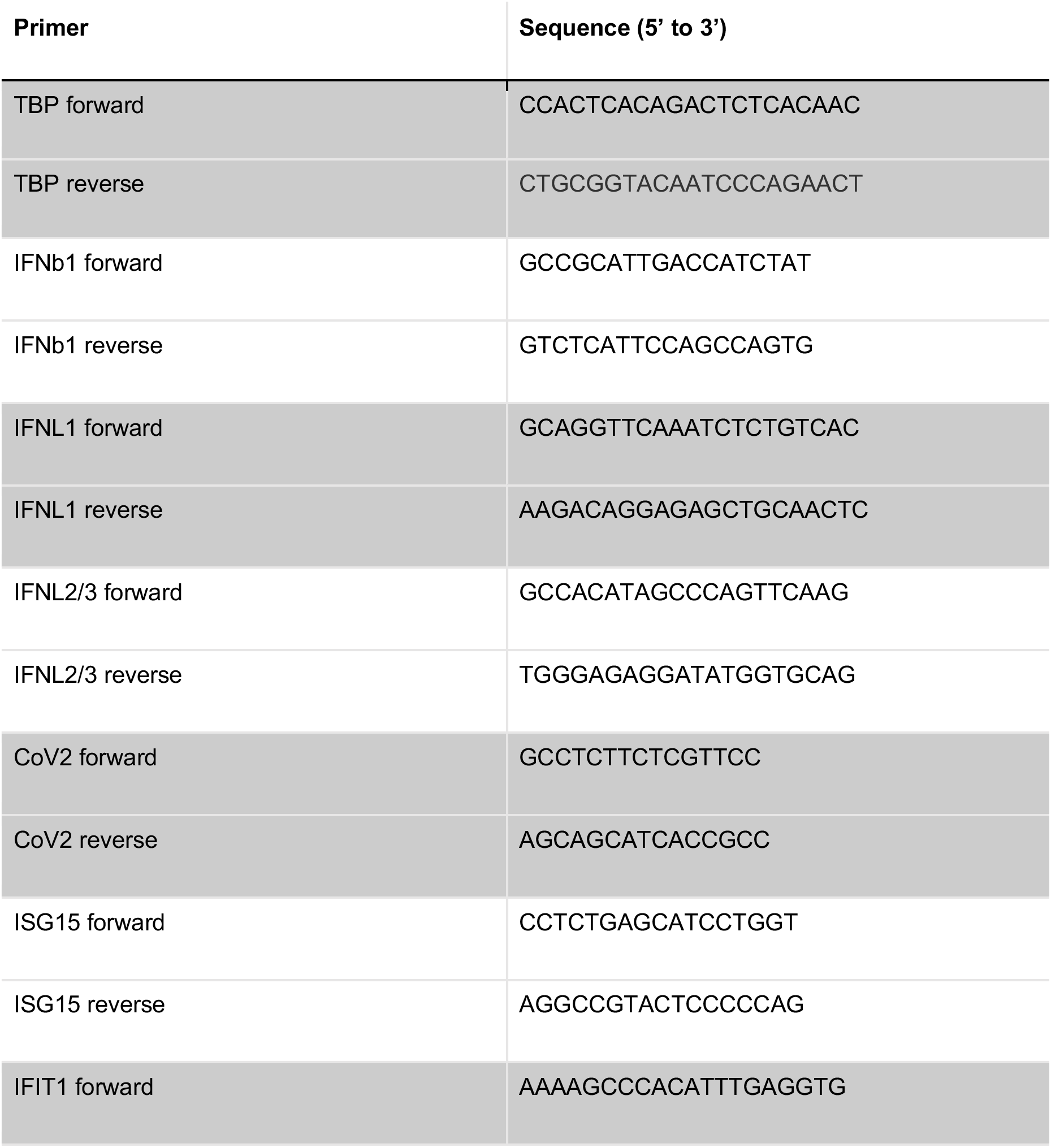
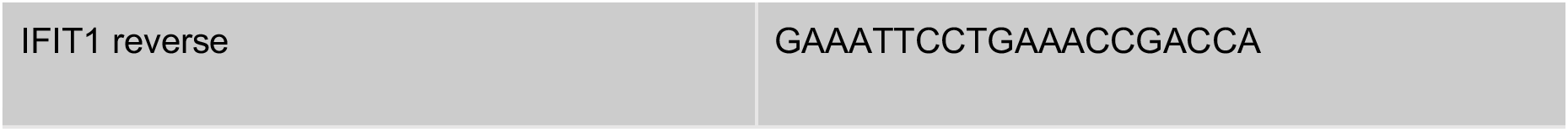

### Indirect immunofluorescence assay

T84 cells were seeded on iBIDI glass bottom 8-well chamber slides 24 h prior to infection. At indicated times post-infection, cells were fixed in 4% paraformaldehyde (PFA) for 20 mins at room temperature (RT). Cells were washed and permeabilized in 0.5% Triton-X for 15 mins at RT. Mouse monoclonal antibody against SARS-CoV NP (Sino biologicals MM05) and mouse monoclonal against J2 (scions) were diluted in phosphate-buffered saline (PBS) at 1/1000 dilution and incubated for 1h at RT. Cells were washed in 1X PBS three times and incubated with secondary antibodies (conjugated with AF488 (Molecular Probes), or AF568 (Molecular Probes) directed against the animal source) and DAPI for 45 mins at RT. Cells were washed in 1X PBS three times and maintained in PBS. Cells were imaged by epifluorescence on a Nikon Eclipse Ti-S (Nikon).

### In-cell western

20,000 Vero E6 cells were seeded per well into a 96-well dish 24 hours prior to infection. 100μL of harvested supernatant was added to the first well. Seven serial 1:10 dilutions were made (all samples were performed in triplicate). Infections were allowed to proceed for 24h. 24 hours post-infection cells were fixed in 2% PFA for 20 mins at RT. PFA was removed and cells were washed twice in 1X PBS and then permeabilized for 10 mins at RT in 0.5% Triton-X. Cells were blocked in a 1:2 dilution of Li-Cor blocking buffer (Li-Cor) for 30 mins at RT. Cells were stained with 1/1000 dilution anti-dsRNA (J2) for 1 h at RT. Cells were washed three times with 0.1% Tween in PBS. Secondary antibody (anti-mouse CW800) and DNA dye Draq5 (Abcam) were diluted 1/10,000 in blocking buffer and incubated for 1 h at RT. Cells were washed three times with 0.1% Tween/PBS. Cells were imaged in 1X PBS on a LICOR imager.

### Organoid dissociation for scRNAseq

2D seeded organoids harvested after 0 (mock), 12 and 24 hours post-infection were washed in cold PBS and incubated in TrypLE Express (Gibco) for 25 min at 37°C. When microscopic examination revealed that cells had reached a single cell state, they were resuspended in DMEM/F12 and spun at 500 xg for 5 mins. Supernatant was removed and the cell pellet was resuspended in PBS supplemented with 0.04% BSA and passed through a 40 μm cell strainer. Resulting cell suspensions were used directly for single-cell RNAseq.

### Single-cell RNA-seq library preparation

Single-cell suspensions were loaded onto the 10x Chromium controller using the 10x Genomics Single Cell 3’ Library Kit NextGem (10x Genomics) according to the manufacturer’s instructions. In summary, cell and bead emulsions were generated, followed by reverse transcription, cDNA amplification (5 μL of of amplified cDNA was set apart for targeted scRNAseq amplification), fragmentation, and ligation with adaptors followed by sample index PCR. Resulting libraries were quality checked by Qubit and Bioanalyzer, pooled and sequenced using HiSeq4000 (Illumina; high-output mode, paired-end 26 × 75 bp).

### Targeted single-cell RNA-sequencing

For targeted scRNAseq, outer and inner primers for targeted amplification were designed using an R package for primer design described in [24] and available through Bioconductor (http://bioconductor.org/packages/release/bioc/html/TAPseq.html). Primers were ordered desalted as ssDNA oligonucleotides and pooled in an equimolar amount, except for the primer targeting SARS-CoV-2 mRNA which was added in 8-fold excess to the outer and inner panel. All primer sequences are described in Supplementary Table 1. Targeted scRNA-seq was performed as previously described [24], except for using amplified cDNA from the 10X Genomics 3’ scRNA-seq protocol as input material. In short, 10 ng of amplified cDNA were used as input for the outer primer PCR and amplified with 10 PCR cycles. A second seminested PCR using 10 ng of Ampure purified outer PCR as input was performed with inner primer mix and 7 cycles of PCR. Then, a third PCR was done adding Illumina adapters. Resulting libraries were quality checked by Qubit and Bioanalyzer, pooled and sequenced using HiSeq4000 (Illumina; high-output mode, paired-end 26 × 75 bp).

### Pre-processing and quality control of scRNAseq data

Raw sequencing data was processed using the CellRanger software (version 3.1.0). Reads were aligned to a custom reference genome created with the reference human genome (GRCh38) and SARS-CoV-2 reference genome (NC_045512.2). The resulting unique molecular identifier (UMI) count matrices were imported into R (version 3.6.2) and processed with the R package Seurat (version 3.1.3). Low-quality cells were removed, based on the following criteria. All cells with mitochondrial reads > 30% were excluded. Second, we limited the acceptable numbers of detected genes. For both types of samples, cells with <1500 or >9000 detected genes were discarded. The remaining data were further processed using Seurat. To account for differences in sequencing depth across cells, UMI counts were normalized and scaled using regularized negative binomial regression as part of the package sctransform. Afterward, ileum and colon organoids samples were integrated independently to minimize the batch and experimental variability effect. Integration was performed using the canonical correlation analysis and mutual nearest neighbor analysis. The resulting corrected counts were used for visualization and clustering downstream analysis and non-integrated counts for any quantitative comparison.

### Pre-processing of targeted scRNAseq

Targeted scRNA-seq pre-processing was done as described in [24]. In summary, following demultiplexing by sample, sequencing data were processed following the workflow provided by Drop-seq tools (v. 1.13, http://mccarrolllab.org/dropseq/) with STAR (v. 2.5.3a) to align reads. To mitigate potential multi-mapping issues, targeted samples were aligned to a custom alignment reference containing only genes of the respective target gene panel, including the SARS-CoV-2 genome. This reference contained the sequences of all target gene loci as individual contigs with overlapping loci merged into one contig. UMI observations were extracted using the Drop-seq tools GatherMolecularBarcodeDistributionByGene program. A custom script (Python v. 3.6.6) was used to filter for chimeric reads with a transcripts-per-transcript (TPT) cutoff of 0.25, and UMI observations were converted to transcript counts. Cell-containing droplets were extracted using the filtered cell barcodes from the scRNASeq data. Other detected cell barcodes droplets were categorized as empty droplets. The infection status for every cell was extracted from the targeted gene expression data by thresholding the SARS-CoV-2 counts using the media expression of the cell containing droplets (Figure S2). Furthermore, Pearson correlation of each targeted gene to its WTA equivalent was calculated.

### Clustering and identification of cell type markers

We performed principal component analysis (PCA) using 3000 highly variable genes (based on average expression and dispersion for each gene). The top 30 principal components were used to construct a shared nearest neighbor (SNN) graph and modularity-based clustering using the Louvain algorithm was performed. Finally, Uniform manifold approximation and projection (UMAP) visualization was calculated using 30 neighboring points for the local approximation of the manifold structure. Marker genes for every cell type were identified by comparing the expression of each gene in a given cluster against the rest of the cells using the receiver operating characteristic (ROC) test. To evaluate which genes classify a cell type, the markers were selected as those with the highest classification power defined by the AUC (area under the ROC curve). These markers along with canonical markers for intestinal and colonic cells were used to annotate each of the clusters of the ileum and colon samples.

### Differential expression analysis

To identify the changes in expression across conditions. Differential expression tests were performed using MAST [47]. To reduce the size of the inference problem and avoid cell proportion bias, separate models were fit for each cell lineage and comparisons between mock, 12 hours, and 24 hours post-infection were performed. False discovery rate (FDR) was calculated by the Benjamini-Hochberg method [48] and significant genes were set as those with FDR of less than 0.05. Subsequently, genes whose mRNAs were found to be differentially expressed were subjected to a gene set overrepresentation analysis using the EnrichR package in R. Furthermore signalling pathways enrichment was calculated using PROGENy.

### Multiplex FISH

Organoids were seeded in expansion medium on glass coverslips. At 24 hours post-seeding, the expansion medium was replaced by differentiation medium and organoids were left to differentiate for 4 days. After validation that organoids were differentiated and contained all expected cell types, organoids were infected, harvested after 12 and 24 hpi, and fixed in 4% PFA for 30 minutes. HiPlex (RNAscope) was performed following the manufacturer's instructions. Briefly, fixed samples were dehydrated with 50%, 70%, 100% ethanol, then treated with protease. All the HiPlex probes were hybridized and amplified together. Probes were designed for genes identified as cell type markers and/or corroborated by literature. The probes used were:

| Gene name | Role of the marker | Catalog number |
| --- | --- | --- |
| ACE2 | SARS-CoV-22 Receptor | 848151 |
| ISG15 | Infection Marker | 311021 |
| nCoV2019-S | SARS-CoV-22 marker | 848561 |

After washing, cell nuclei were counterstained with DAPI, and samples were mounted using ProLong Gold Antifade Mountant. Imaging was performed with the camera Nikon DS-Qi2 (Nikon Instruments) with the Plan Fluor 20x objective (Nikon Instruments) mounted on the Nikon Ti-E inverted microscope (Nikon Instruments) in bright-field and fluorescence (DAPI, GFP, Cy3 and Cy5 channels). The microscope was controlled using the Nikon NIS Elements software. After each round, fluorophores were cleaved and samples moved on to the next round of the fluorophore detection procedures. All images from all rounds of staining were then registered with each other to generate images using HiPlex image registration software (ACD Bio). Further brightness and contrast adjustments were performed using Fiji.

The HiPlex probe fluorescent signal was used to determine the ACE2 and ISG15 RNA expression levels, as well as the SARS-CoV-2 infection levels. To obtain a resolution at a single cell level, first nuclei segmentation and classification was done on raw DAPI images using the Pixel Classification + Object Classification workflow from ilastik 1.2.0. The resulting Object Prediction masks represented all nuclei as individual objects in a 2D plane and were saved as 16bit Tagged Image File Format. To measure the single cell fluorescent intensity for the ACE2, ISG15 and SARS-CoV-2 probes, a pipeline using CellProfiler 3.1.9 was developed. Briefly, first the raw grayscale images corresponding to the ACE2, ISG15 and SARS-CoV-2 probe fluorescent signals were uploaded on the pipeline. These images were specified as images to be measured. The corresponding Object Prediction masks previously generated by ilastik were then uploaded, converted into binary nuclei masks and used to define the objects to be measured. Finally, with a MeasureObjectIntesity module the fluorescence intensity features, the cell number and the single cell localization were measured for the identified objects from the binary nuclei mask. The outcome was exported to a spreadsheet and contained the localization as well as the mean intensity units rescaled from 0 to 1 of ACE2, ISG15 and SARS-CoV-2 fluorescent signals for each single cell.

To determine the infection status for every cell a threshold was calculated using the SARS-CoV-2 mean fluorescent intensity signal of mock treated versus representative infected cells. The threshold was set to 0.015 mean intensity units. Due to the probe quality, the ACE2 fluorescent signal showed strong variations between different images and hence technical replicates. To minimize the variability, for each individual image the ACE2 mean intensity signal was normalized and rescaled from 0 to 1, 0 corresponding to the lowest and 1 to the highest ACE2 mean intensity signal of a cell from the corresponding image. Finally, the SARS-CoV2 mean intensity signal was plotted against the normalized ACE mean intensity signal or the ISG15 mean intensity signal using GraphPad Prism Version 6.0.

### Transcription factor activity along diffusion map pseudotime

Raw counts were normalised using the SCTransform method implemented in Seurat v. 3.2.0, regressed out over UMI counts. Transcription factor activities were then calculated using pySCENIC v 0.10.13 (Sande et al. 2020). Independently, the diffusion maps were computed using the destiny R library v 3.2.0. For the inference of pseudotime in Ileum a set of curated genes related to IFN signaling from Reactome were used as input. For colon, we also added genes with highest variability (standard deviation > 2). Visualisation of the TF activities along trajectories were carried out with custom R scripts (v 4.0.2).

## Data and Code Availability

The raw sequencing generated during this study is available at the NCBI Gene Expression Omnibus (accession no. GSE156760). The authors declare that all other data supporting the findings of this study are within the manuscript and its supplementary files.

