## Supplementary Data for "Single-cell analyses reveal SARS-CoV-2 interference with intrinsic immune response in the human gut"

**This PDF file includes:**

Fig S1-S10
Table S1

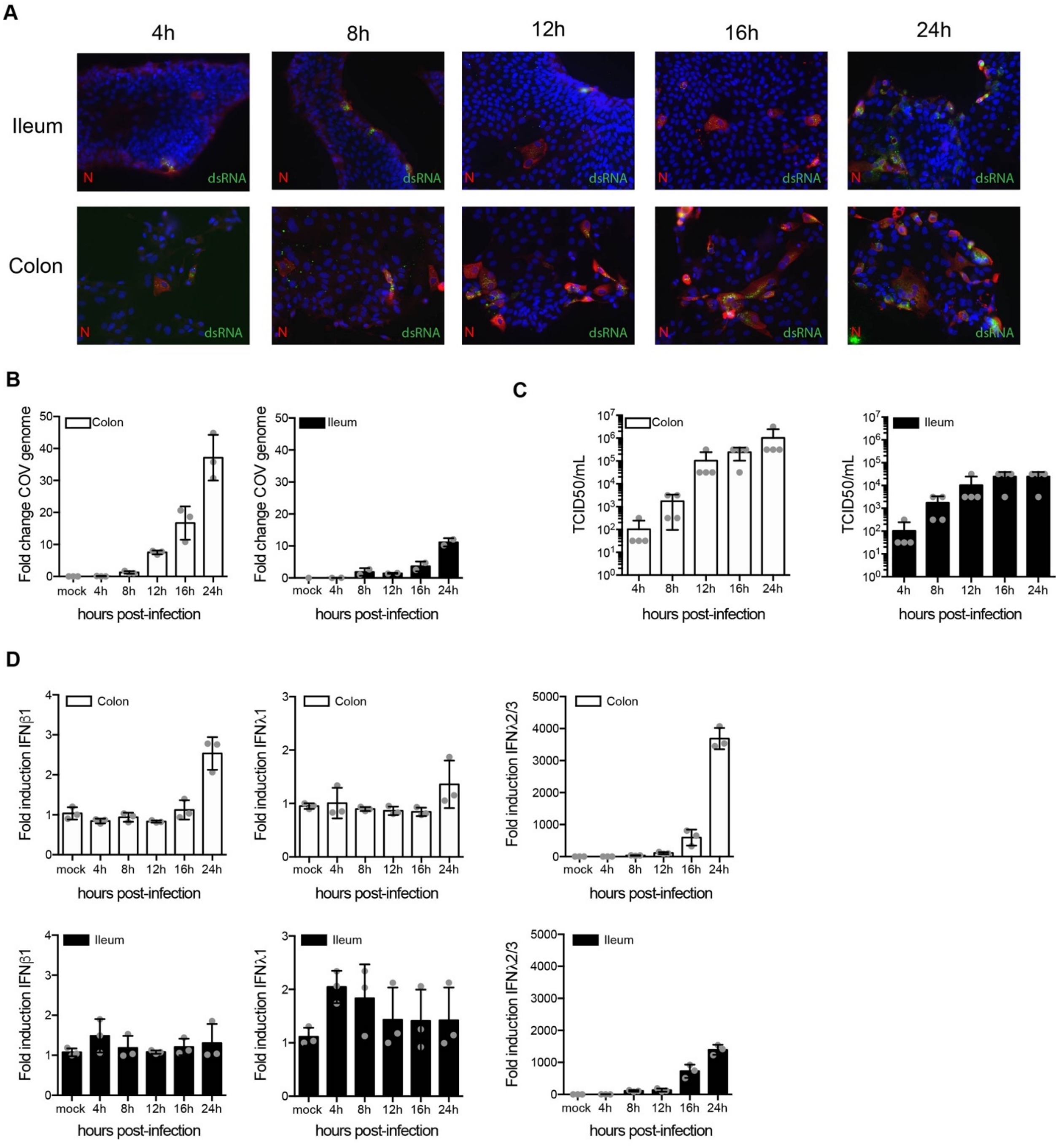


**Fig. S1.**

**Ileum- and colon-derived organoids support SARS-CoV-2 replication and spread.** Ileum- and colon-derived organoids were seeded in 2D and were infected with SARS-CoV-2. A. At 4, 8, 12,

16 and 24 hpi, samples were fixed and analyzed by immunofluorescence for the SARS-CoV-2 N protein (red), dsRNA (green) and nuclei were stained with DAPI (blue). Three biological replicates were performed, representative images are shown. B. At indicated times, RNA was harvested and the amount of virus replication was monitored by q-RT-PCR. Three biological replicates were performed. Error bar indicates standard deviation. C. At indicated time points, supernatants were collected and the amounts of infectious viruses produced de novo were titrated on naïve Vero cells. Four biological replicates were performed. Error bar indicates standard deviation. D. At indicated time points, RNA was harvested and the upregulation of IFNβ1, INFλ1 and IFNλ2/3 was analyzed by q-RT-PCR. Three biological replicates were performed. Error bar indicates standard deviation.

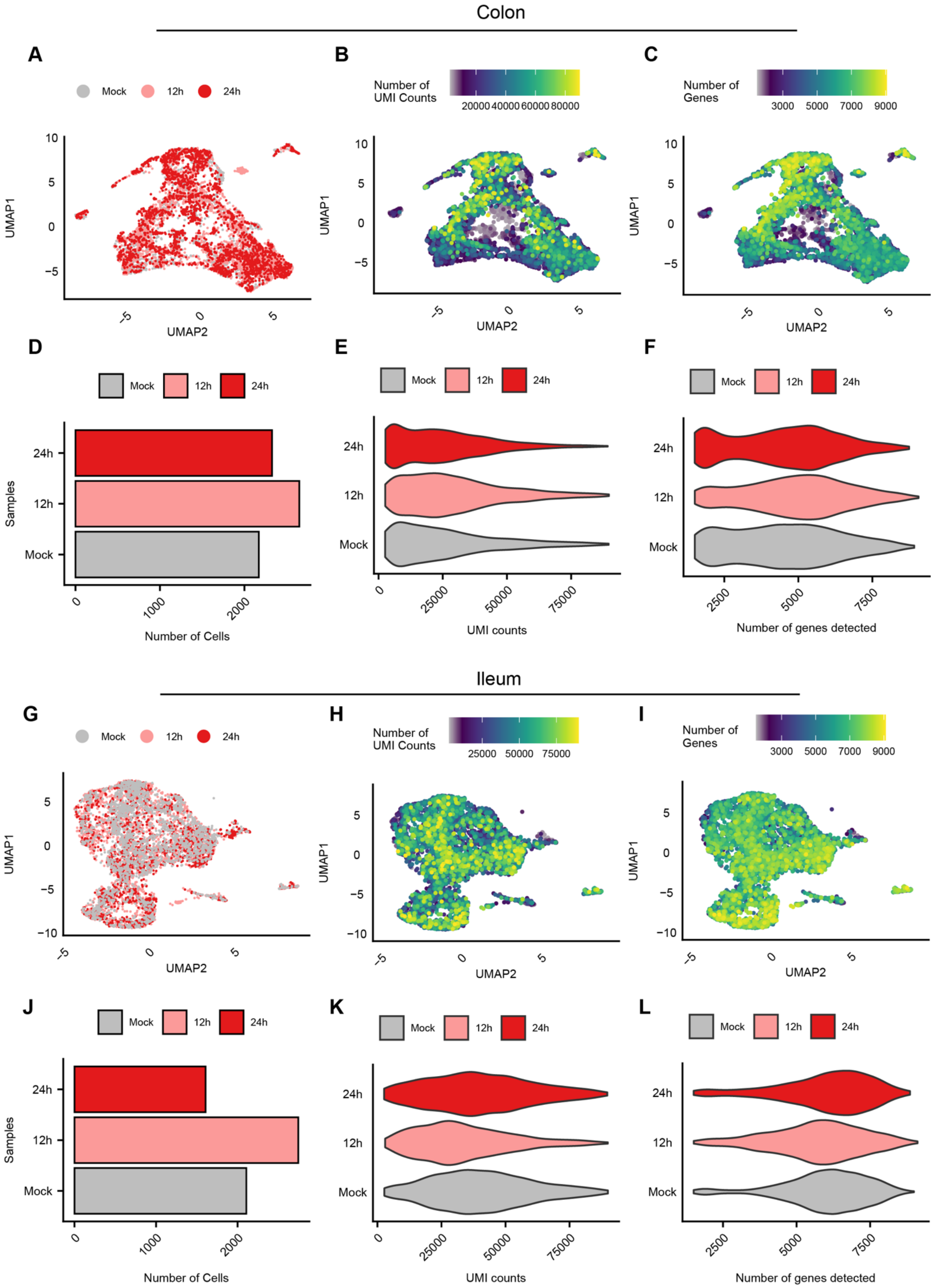

Colon
A
Mock 12h 24h
UMAP1
UMAP2
B
Number of UMI Counts
20000 40000 60000 80000
C
Number of Genes
3000 5000 7000 9000
D
Mock 12h 24h
Samples
Number of Cells
E
UMI counts
F
Number of genes detected
Ileum
G
H
Number of UMI Counts
25000 50000 75000
I
Number of Genes
J
K
L

**Fig. S2.**

**General information of single-cell RNA-seq. A-F**. Data correspond to scRNAseq of mock and SARS-CoV-2 infected colon organoids. **G-L**. Data correspond to scRNAseq of mock and SARS-CoV-2 infected ileum organoids. **A.** and **G.** Uniform manifold approximation and projection (UMAP) embedding of scRNA-Seq data from mock and SARS-CoV-2 infected organoids at 12 and 24 hpi. Conditions are color coded. **B.** and **H.** Number of UMI counts. **C.** and **I.** Number of genes sequenced in each cell **D.** and **J.** Number of cells in scRNAseq datasets for each condition (mock, 12 and 24 hpi). **E.** and **K.** Violin plots depicting the number of UMI per cell for each condition (mock, 12 and 24 hpi). **F.** and **L.** Violin plots depicting the number of genes detected per cell for each condition (mock, 12 and 24 hpi).

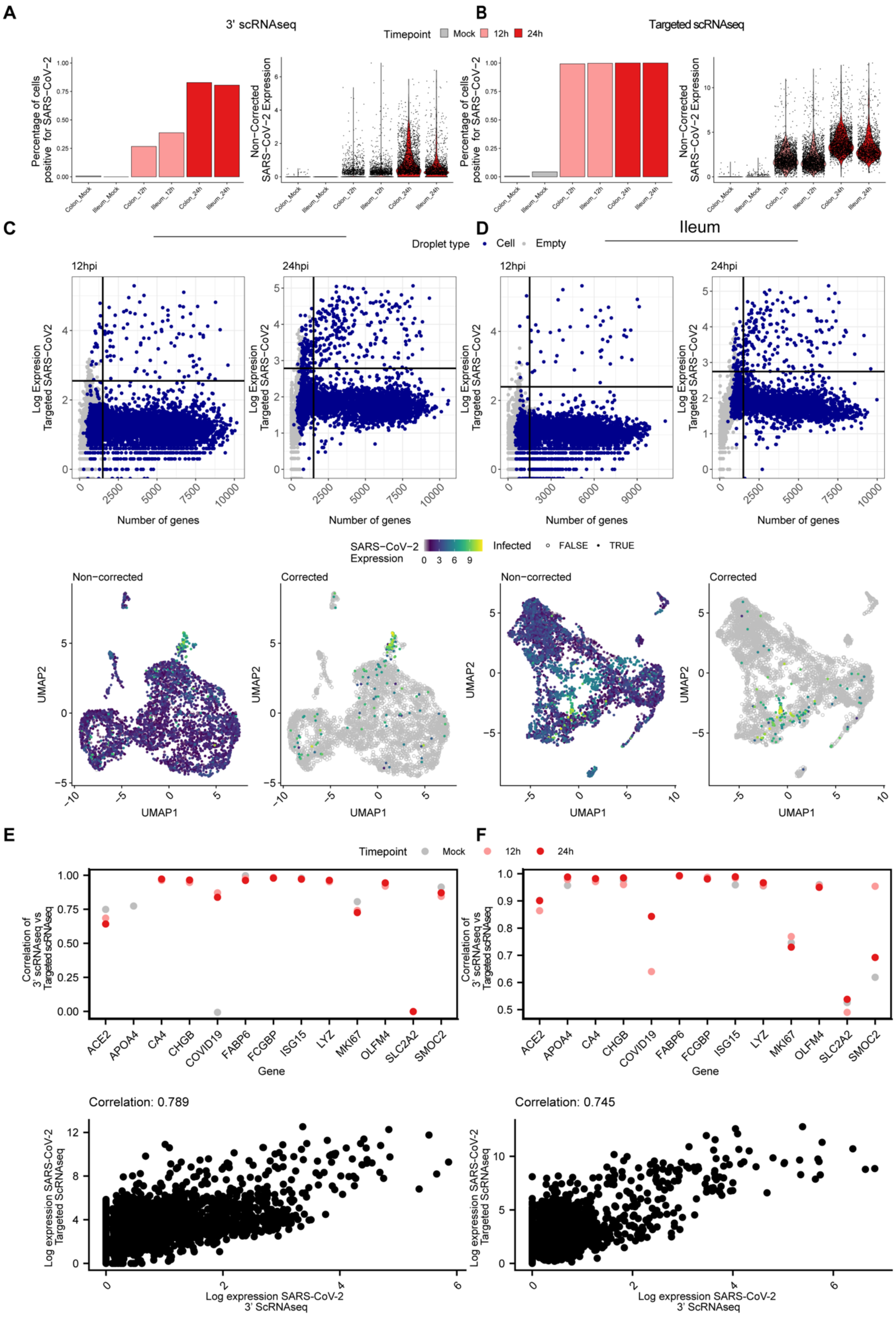

A
3' scRNAseq
B
Targeted scRNAseq
Timepoint Mock 12h 24h
Percentage of cells positive for SARS-CoV-2
Non-Corrected SARS-CoV-2 Expression
C
D
Ileum
Droplet type Cell Empty
12hpi
24hpi
Log Expression Targeted SARS-CoV2
Number of genes
SARS-CoV-2 Expression
Infected FALSE TRUE
Non-corrected
Corrected
UMAP1
UMAP2
E
F
Correlation of 3' scRNAseq vs Targeted scRNAseq
Gene
ACE2
APOA4
CA4
CHGB
COVID19
FABP6
FCGBP
ISG15
LYZ
MKI67
OLFM4
SLC2A2
SMOC2
Correlation: 0.789
Correlation: 0.745
Log expression SARS-CoV-2 Targeted ScRNAseq
Log expression SARS-CoV-2 3' ScRNAseq

**Fig. S3.**

**Identification of SARS-CoV-2 infected cells using targeted scRNAseq datasets. A-B.** Violin plots displaying SARS-CoV-2 expression for the 10x genomics 3' scRNAseq and for the targeted scRNAseq for mock-infected and SARS-CoV-2 infected colon and ileum organoid at 12 and 24 hpi. **C.** (Top panels) SARS-CoV-2 expression as function of the number of genes per droplet from the targeted scRNAseq datasets of colon organoids at 12 and 24 hpi. Droplet types are colored for droplets containing cells and empty droplets. The vertical line represents the threshold for droplets containing <2000 genes per cell. The horizontal line represents the threshold used to define the baseline of infection at 12 and 24 hpi. (Bottom panels) Uniform manifold approximation and projection (UMAP) embedding of the scRNA-Seq data of infected colon organoids depicting SARS-CoV-2 infected cells. (Left) Expression of SARS-CoV-2 in non-corrected datasets. (Right) Expression of SARS-CoV-2 in corrected datasets using the thresholds determined in top panel dotplots. **D.** Same as C. but for ileum organoids. **E-F** (Top panels) Correlation of the expression of each gene in 10x genomics 3' scRNAseq against their expression in targeted scRNAseq across conditions. (Bottom panels) SARS-CoV-2 expression in each infected cell in 10x genomics 3' scRNAseq against targeted scRNAseq. Colon (left) and Ileum (right).

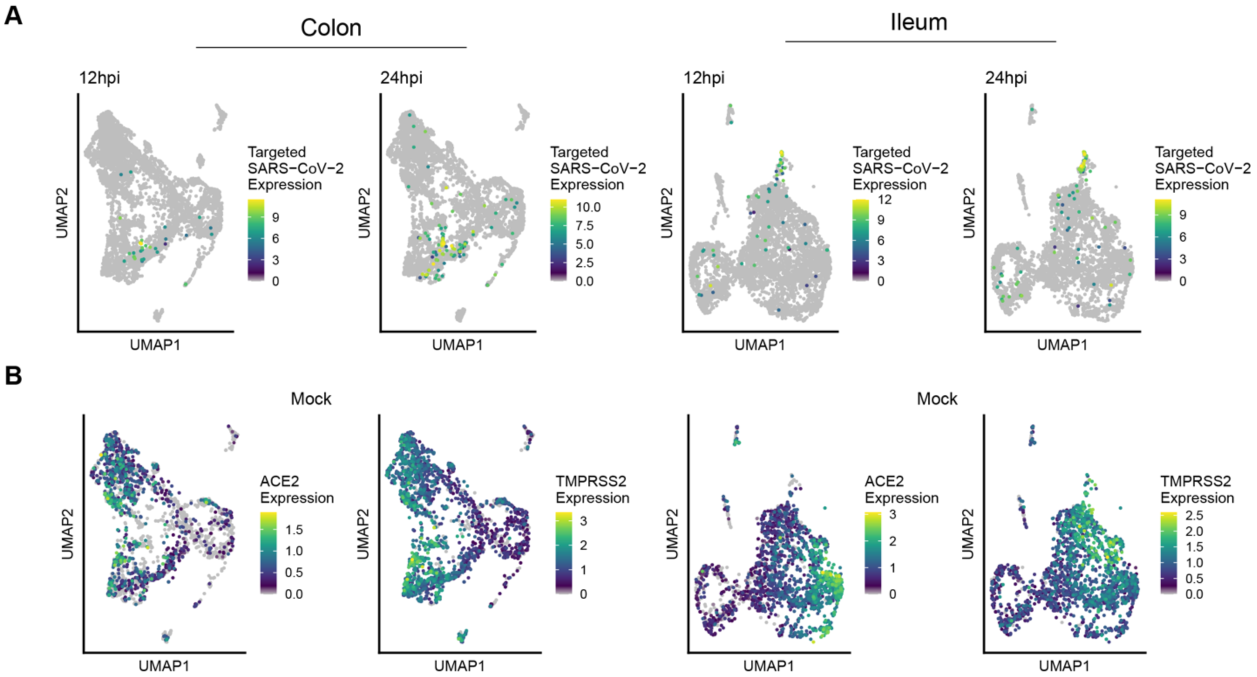


**Figure S4. Expression of SARS-CoV-2 and ACE2 in colon- and ileum-derived organoids. A.** Uniform manifold approximation and projection (UMAP) embedding of the scRNA-Seq data of infected colon organoids at 12 and 24 hpi, coloured by the corrected targeted normalized expression of SAR-Cov-2. Colon (left) and Ileum (right). **B.** Same as A. but for ACE2 and TMPRSS2 expression at mock .

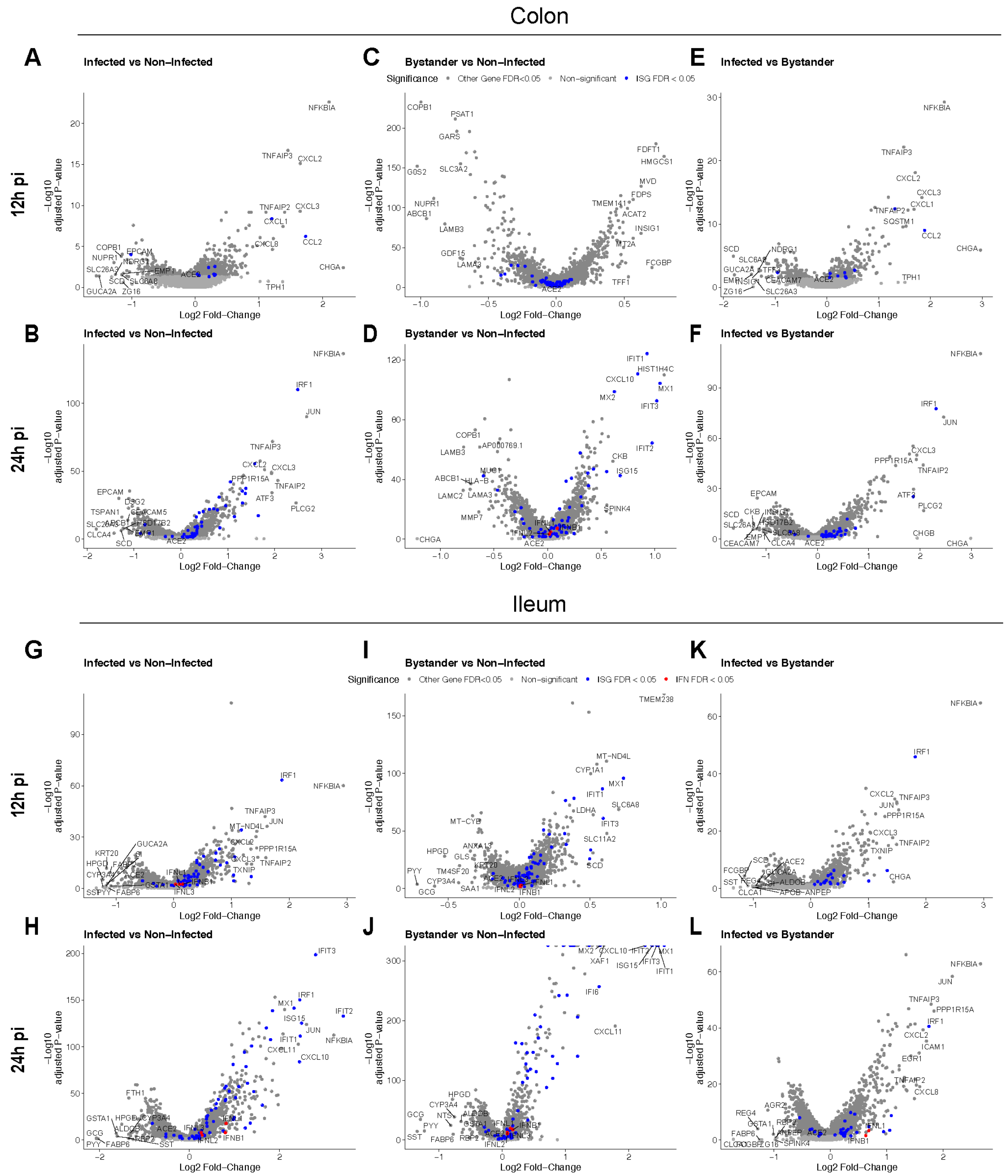


**Figure S5. Differential response of infected and bystander cells to SARS-CoV-2 infection.** **A-F**. Volcano plots displaying the genes that are differentially expressed upon SARS-CoV-2 infection of colon organoids. **A-B.** infected *vs*. mock infected cells at 12 hpi (A) and 24 hpi (B), **C-D.** bystander *vs.* mock infected cells at 12 hpi (C) and 24 hpi (D), and **E-F.** infected *vs.* bystander cells at 12 hpi (E) and 24 hpi (F). The statistical significance (-log10 adjusted p-value) is shown as a function of the log2 fold change. **G-L**. Same as A-F. but for ileum organoids.

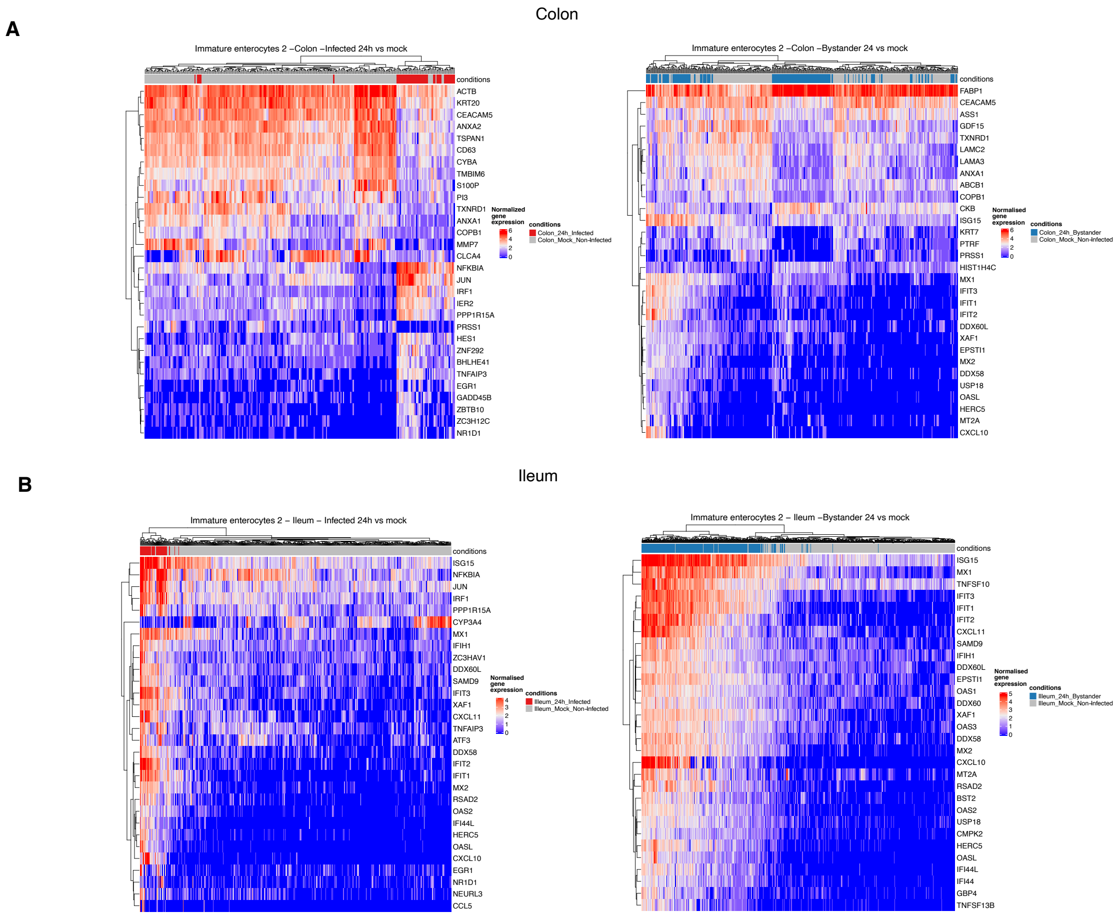


**Figure S6. Gene expression profiles showing the top 30 genes with highest fold change in the differential expression analysis. A.** Gene expression profiles of immature enterocytes 2 subpopulation in colon-derived organoids 24 hpi**.** (Left) Infected *vs*. mock infected cells at 24 hpi and (right) bystander *vs.* mock infected cells. **B.** Same as A but for ileum-derived organoids.

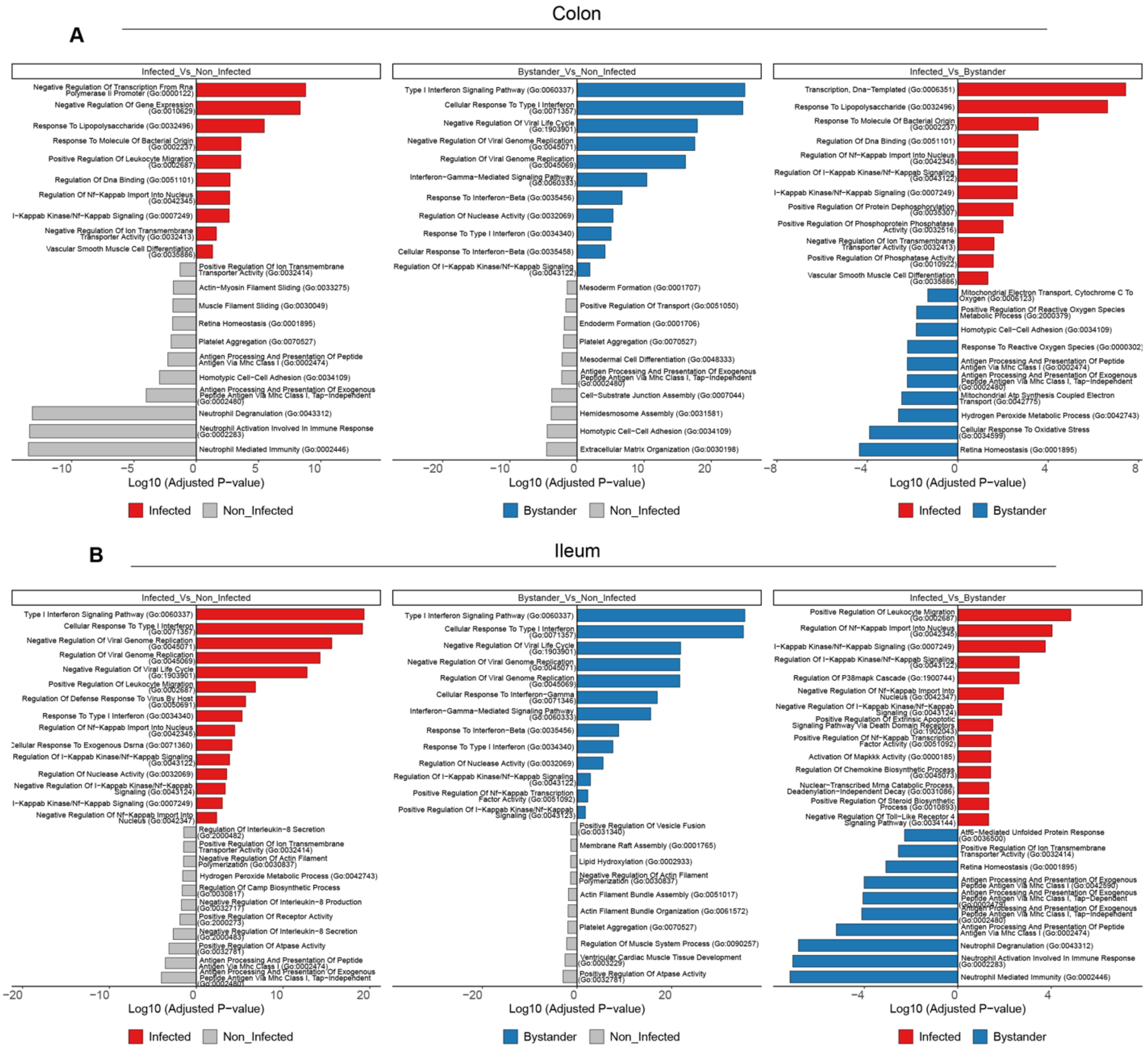


**Figure S7. Gene enrichment analysis in infected and bystander cells upon SARS-CoV-2 infection. A.** Gene Ontology (GO) enrichment analysis was performed on the genes that are differentially expressed (FDR<0.05) upon SARS-CoV-2 infection of colon organoids in infected relative to mock cells (left panel), in bystander relative to mock cells (middle panel) and in infected relative to bystander cells (right panel). **B.** same as A. but for ileum organoids.

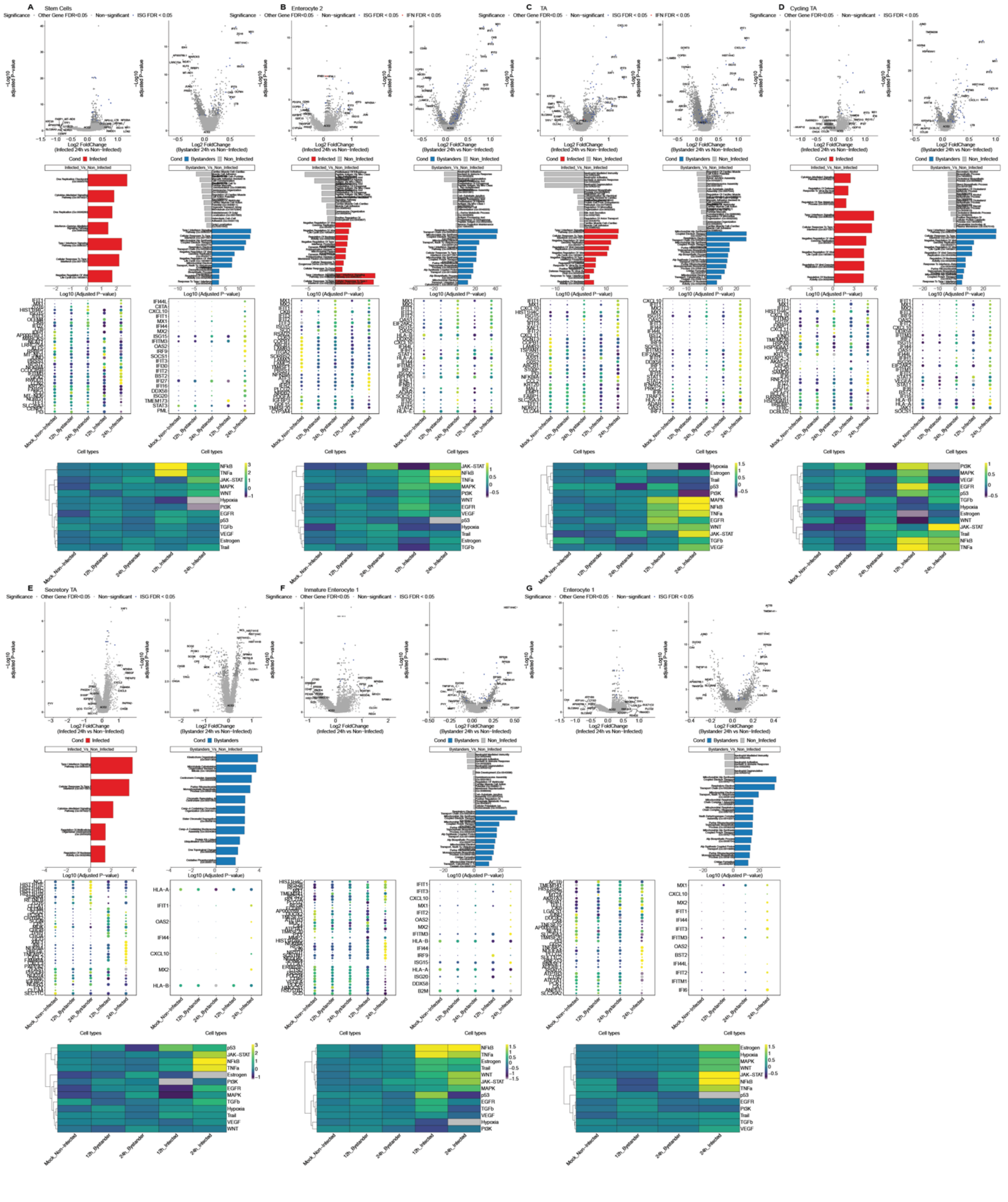


**Figure S8. Cell type specific differential response of infected *vs.* bystander cells upon SARS-CoV-2 infection of colon-derived organoids.** (top two panels) Volcano plots displaying the genes that are differentially expressed in infected and bystander cells relative to mock-infected cells. The statistical significance (-log10 adjusted p-value) is shown as a function of the log2 fold change. (upper middle panels) Gene Ontology (GO) enrichment analysis was performed on the genes that are differentially expressed (FDR<0.05) upon SARS-CoV-2 infection of colon organoids in infected relative to mock-infected cells (left panel) and in bystander relative to mock-infected cells (right panel). (lower middle panels, left) Dot plot of the top most differentially expressed genes upon SARS-CoV-2 infection in mock, infected and bystander cells at 12 and 24 hpi. The dot size represents the percentage of cells expressing the gene; the color represents the average expression across the cell type. (lower middle panels, right) Dot plot of the top most differentially expressed ISGs upon SARS-CoV-2 infection in  mock, infected and bystander cells at 12 and 24 hpi. The dot size represents the percentage of cells expressing the gene; the color represents the average expression across the cell type. (lower panel) Heatmap of signaling pathway enrichment inferred by PROGENy for the given cell type. **A.** Stem cells, **B**. Enterocyte 2. **C**. TA cells, **D.** Cycling TA, **E.** Secretory TA, F. Immature enterocyte 1 and **G.** Enterocyte 1.

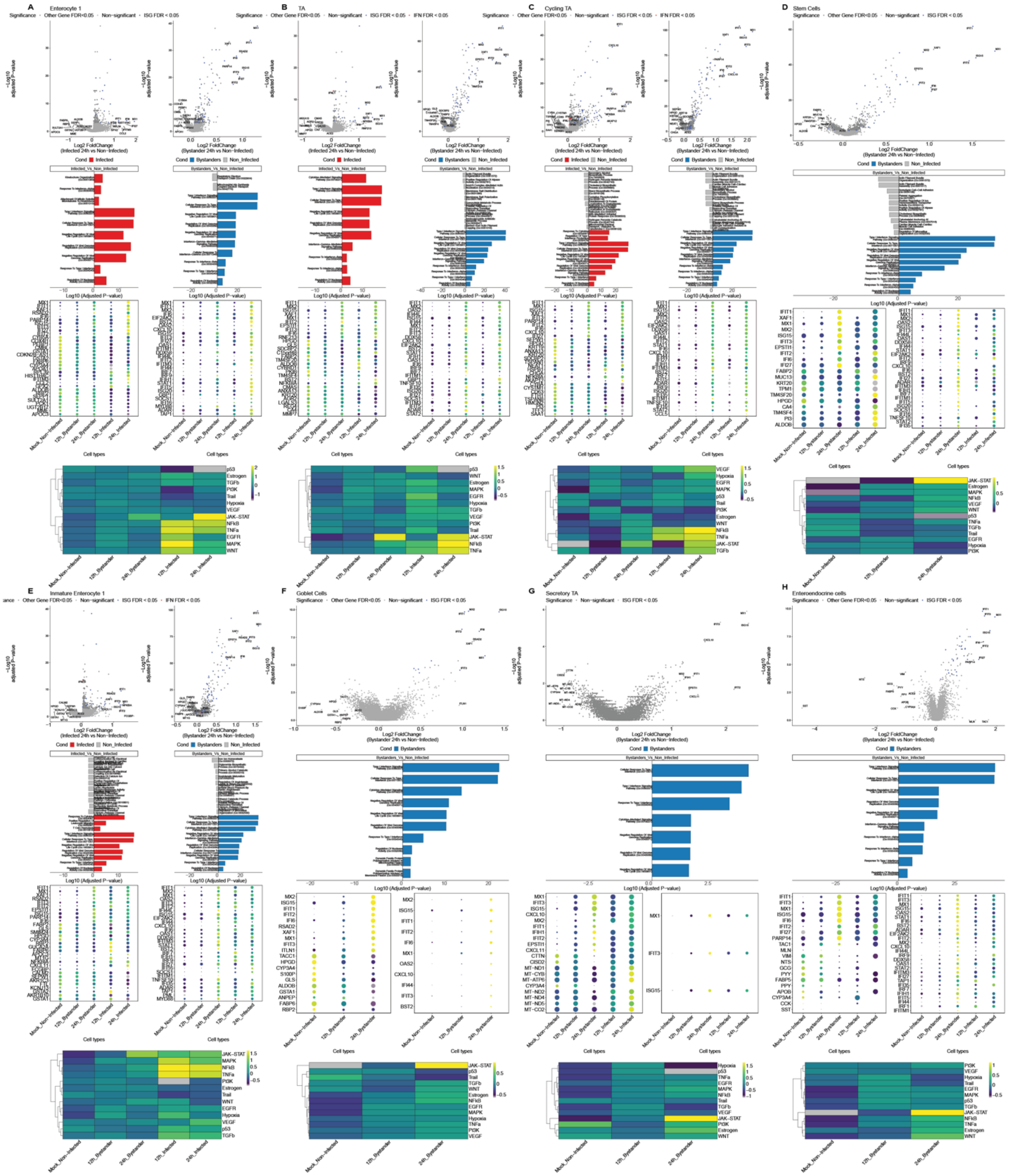

**Figure S9. Cell type specific differential response of infected *vs.* bystander cells upon SARS-CoV-2 infection of ileum-derived organoids.** Same as figure S8 but for ileum organoids. **A.** Enterocyte 1, **B.** TA cells, **C.** Cycling TA, **D.** Stem cells, **E.** Immature enterocyte 1, **F.** Goblet cells and **G.** Secretory TA.

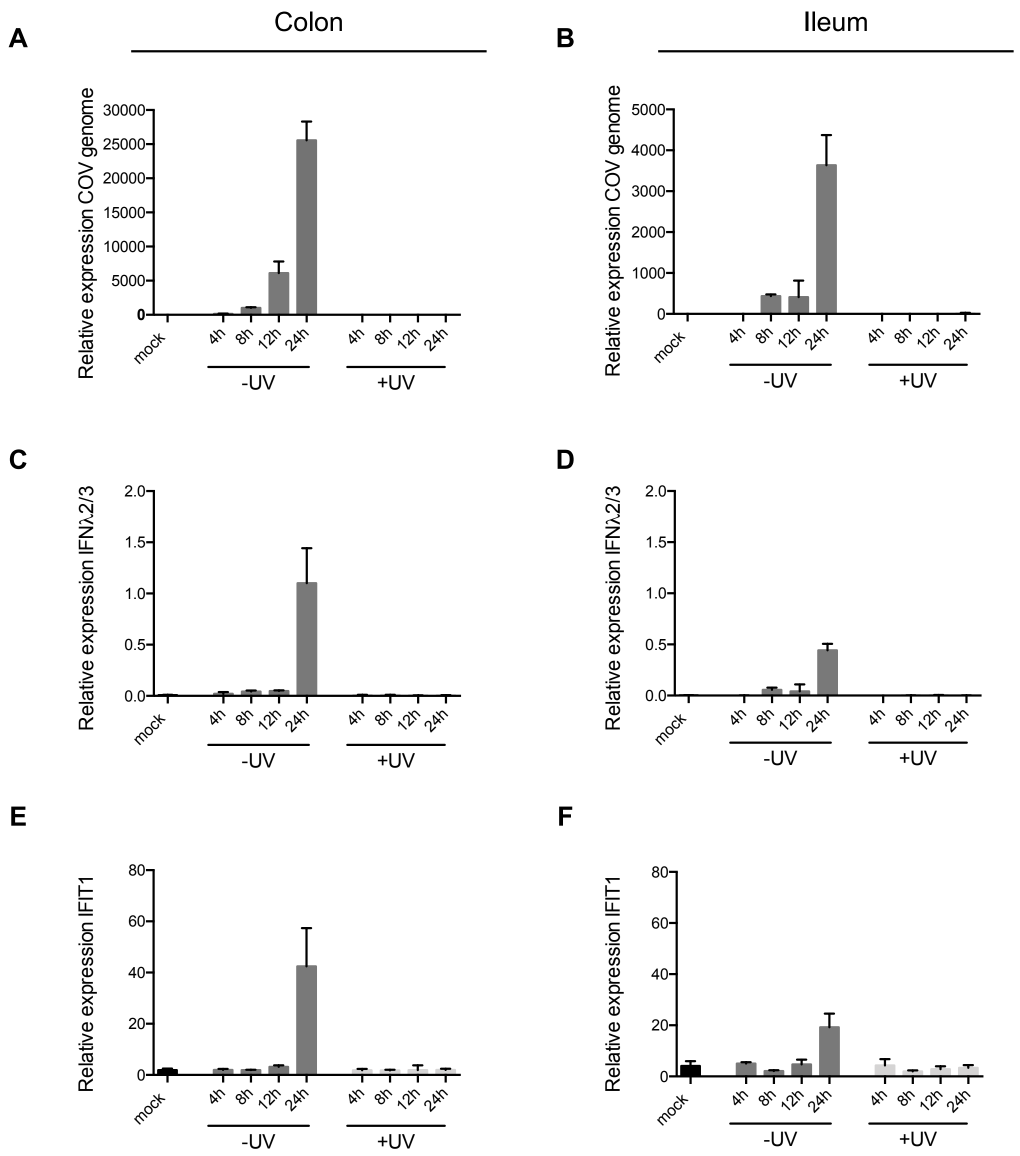


**Figure S10. Interferon induction requires virus replication.** Ileum- and colon-derived organoids were seeded in 2D and were infected with live SARS-CoV-2 (-UV) or UV-inactivated SARS-CoV-2 (+UV). At 4, 8, 12 and 24 hpi, RNA was harvested and analyzed for **A-B.** replication of SARS-CoV-2 genome, **C-D.** induction of type III interferon (IFNλ2/3) and **E-F.** induction of the interferon-stimulated gene IFIT1. Three biological replicates were performed for each experiment. Error bars indicate standard deviation.

| Outer primer | Sequence | Inner primer | Sequence |
|---|---|---|---|
| COVID19_outer | ACCACACAAGGCAGATGGGC | COVID19_inner | GTCTCGTGGGCTCGGAGATGTGTATAAGAGACAGCATTTTCACCGAGGCCACGC |
| ACE2_outer | CTGGGAACTGGTGTAGCTGCA | ACE2_inner | GTCTCGTGGGCTCGGAGATGTGTATAAGAGACAGTCCAGGGAACAGGTAGAGGAC |
| APOA4_outer | CGAGGGGCTGCAGAAGTCAC | APOA4_inner | GTCTCGTGGGCTCGGAGATGTGTATAAGAGACAGGGCCACTTGAGCTTCCTGGAG |
| CHGB_outer | TCTGAGGAGCCGGTGAGCAC | CHGB_inner | GTCTCGTGGGCTCGGAGATGTGTATAAGAGACAGCTGTCATTGGAGCGGTGGGC |
| FABP6_outer | AGCATGGCTTTCACCGGCAA | FABP6_inner | GTCTCGTGGGCTCGGAGATGTGTATAAGAGACAGTGGTCCCAGCACTACTCCGG |
| FCGBP_outer | CCTGGTACCGTGTAGTTGCCG | FCGBP_inner | GTCTCGTGGGCTCGGAGATGTGTATAAGAGACAGCTCCCTGCTAGTCCGCCAGA |
| ISG15_outer | GCAGCTCCATGTCGGTGTCA | ISG15_inner | GTCTCGTGGGCTCGGAGATGTGTATAAGAGACAGGCTGGTGGTGGACAAATGCG |
| LYZ_outer | GCCCGGCCACATTCAGTTCT | LYZ_inner | GTCTCGTGGGCTCGGAGATGTGTATAAGAGACAGGGCAAAATACCAGCTGATGAAGGC |
| MKI67_outer | GCTCTGCTCCCGCCTGTTTT | MKI67_inner | GTCTCGTGGGCTCGGAGATGTGTATAAGAGACAGTCCACTTTGCCCCCTGTCCT |
| OLFM4_outer | ACCAACCCCCTTCTACTGCCT | OLFM4_inner | GTCTCGTGGGCTCGGAGATGTGTATAAGAGACAGTGCCTTTGTTTAAGCCTGGAACT |
| SLC2A2_outer | TGGCTAGTGGCAATAAGTTCCA | SLC2A2_inner | GTCTCGTGGGCTCGGAGATGTGTATAAGAGACAGTCACCTGATCATATAGCGTGGGT |
| SMOC2_outer | TTCATCCGAGTGGCACTGGC | SMOC2_inner | GTCTCGTGGGCTCGGAGATGTGTATAAGAGACAGGGCCCCAAATCCCCTGAGAC |
| SST_outer | CTGCTGATCCGCGCCTAGAG | SST_inner | GTCTCGTGGGCTCGGAGATGTGTATAAGAGACAGAGCTGCTGTCTGAACCCAACC |

Table S1. Primers used for targeted sc-RNA seq
